# Phylogenetic modeling of enhancer shifts in African mole-rats reveals regulatory changes associated with tissue-specific traits

**DOI:** 10.1101/2023.01.10.523217

**Authors:** Elise Parey, Stephanie Frost, Ainhoa Uribarren, Thomas J. Park, Markus Zoettl, Ewan St. John Smith, Camille Berthelot, Diego Villar

## Abstract

Changes in gene regulation have long been thought to underlie most phenotypic differences between species. Subterranean rodents, and in particular the naked mole-rat, have attracted substantial attention due to their proposed phenotypic adaptations, which include hypoxia tolerance, metabolic changes and cancer resistance. However, it is largely unknown what regulatory changes may associate with these phenotypic traits, and whether these are unique to the naked mole-rat, the mole-rat clade or also present in other mammals. Here, we investigate regulatory evolution in heart and liver from two African mole-rat species and two rodent outgroups using genome-wide epigenomic profiling.

First, we adapted and applied a phylogenetic modeling approach to quantitatively compare epigenomic signals at orthologous regulatory elements, and identified thousands of promoter and enhancer regions with differential epigenomic activity in mole-rats. These elements associate with known mole-rat adaptation in metabolic and functional pathways, and suggest candidate genetic loci that may underlie mole-rat innovations. Second, we evaluated ancestral and species-specific regulatory changes in the study phylogeny, and report several candidate pathways experiencing stepwise remodeling during the evolution of mole-rats – such as the insulin and hypoxia response pathways. Third, we report non-orthologous regulatory elements overlap with lineage-specific repetitive elements and appear to modify metabolic pathways by rewiring of HNF4 and RAR/RXR transcription factor binding sites in mole-rats.

These comparative analyses reveal how mole-rat regulatory evolution informs previously reported phenotypic adaptations. Moreover, the phylogenetic modeling framework we propose here improves upon the state-of-the-art by addressing known limitations of inter-species comparisons of epigenomic profiles, and has broad implications in the field of comparative functional genomics.

## INTRODUCTION

Most phenotypic changes across mammals are thought to arise from differences in gene regulation. African mole-rats are a group of rodents displaying unusual longevity [1] and evolutionary adaptations to their subterranean environment (reviewed in [2, 3]), including cooperative behaviour [4, 5], resistance to hypoxia [6–8], anoxia [9] and hypercapnia [8], metabolic adaptations [9, 10] and pain insensitivity [11]. These unusual traits have prompted genome sequencing of both the naked mole-rat [12] (*Heterocephalus glaber*) and the Damaraland mole-rat [13] (*Fukomys damarensis*), and genomic investigations on species-specific changes in protein sequences and signatures of positive selection [14, 15]. Recent work has applied proteomic [16] and metabolomic [10, 17] approaches to the study of molerat traits, yet the extent to which mole-rat specific changes in gene regulation may underlie these adaptations remains unexplored. Moreover, the significance and uniqueness of molerat phenotypic traits has been subject to debate [18], in part due to many observations being limited to comparisons between naked mole-rat and mouse.

Mammalian gene regulation is largely enacted by collections of non-coding promoter and enhancer regions, known to bind hundreds of transcription factors combinatorially [19–21]. Previous studies have extensively documented the evolution of mammalian regulatory elements [22–26], which is especially fast for enhancers. Comparative analyses across species, tissues and developmental stages have suggested a greater functional relevance for conserved regulatory activity [27–30]. In contrast, lineage-specific elements appear partly compensatory of proximally lost events [27, 31] and typically arise in genomic regions with pre-existing regulatory activity [22, 32]. Although mammalian adaptations in gene regulation have been linked to the lineage-specific expansion of repetitive elements [33–35], identifying the subsets of rapidly-evolving non-coding elements associated with lineage-specific shifts in regulatory activity has proved more challenging – in part due to limitations in comparative approaches for functional genomics data [36, 37].

Here, we applied a phylogeny-aware approach to quantitatively compare epigenomically-defined promoters and enhancers across two tissues from four rodents, including two molerat species and two rodent outgroups. Our results identify widespread mole-rat specific shifts in promoter and enhancer activities, and inform the gene regulatory component of several mole-rat adaptations. Lastly, we investigate the contribution of repetitive elements to lineagespecific gene regulation, and report that SINE repeats provide new transcription factor binding sites associated with mole-rat modifications in liver metabolism.

## RESULTS

### Comparative epigenomics of liver and heart regulatory activities in mole-rats

To investigate the contribution of enhancer evolution to tissue-specific gene regulation in molerats, we generated histone mark ChIP-sequencing data to identify promoters and enhancers active in two somatic tissues with distinct metabolic and functional roles (liver and heart) from naked mole-rat, Damaraland mole-rat, and two outgroup rodents (guinea pig and mouse; Figure 1A, Figure S1 and Table S1). Using two to six replicates for each combination of species, histone mark and tissue (Table S1), we obtained an average of around 20,000 promoters and 50,000 enhancers reproducibly detected across individuals (Methods, Figure 1B). In the four study species, promoters show over 80% commonality across tissues (Figure 1C, Figure S1), in line with their general association with gene expression [23, 25]. In contrast, liver and heart enhancers in each of the four rodent species are largely distinct, with just under 30% being typically common to both tissues (Figure 1C, Figure S1), consistent with their role as tissue-specific cis-regulators [29, 33].

**Figure 1:**
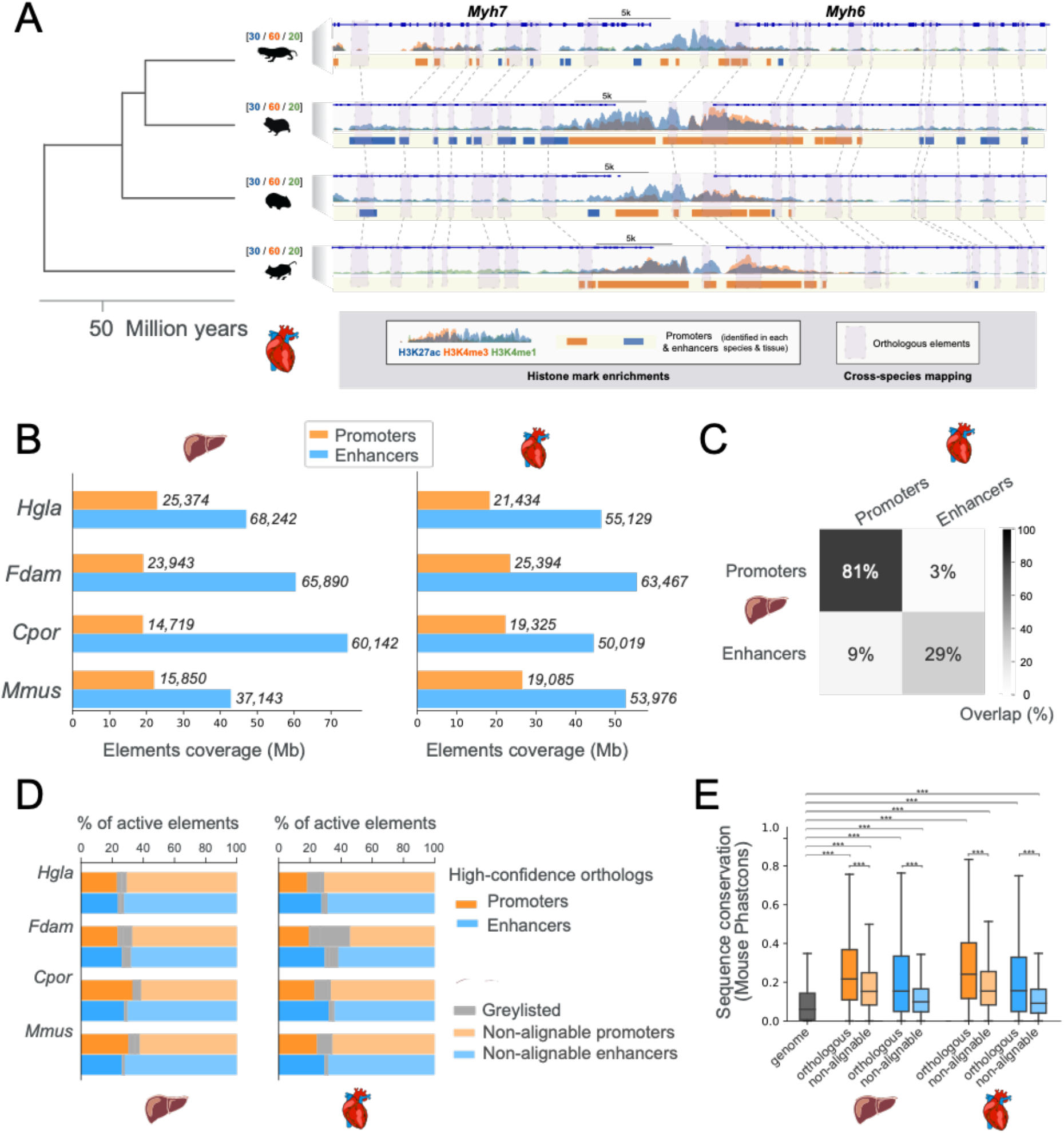
Comparative epigenomics of mole-rat promoters and enhancers in heart and liver. **A.** Epigenomic profiling and cross-mapping approach exemplified by the Myh6/Myh7 heart locus. H3K27ac (blue), H3K4me3 (orange) and H3K4me1 (green) histone marks enrichment in heart are shown for each of the four species (Hgla: naked-mole rat, Fdam: Damaraland mole-rat, Cpor: guinea pig, Mmus: mouse), with scales above species names indicating fold enrichment over input (averaged across replicates). Identified promoters and enhancers are represented by orange and blue boxes, respectively. Purple boxes connected by dashed lines correspond to orthologous promoters and enhancers in each species. **B.** Coverage and numbers of promoters and enhancers by species and tissue. Bars correspond to total genomic coverage, number of elements are indicated at the right of bars. **C.** Overlap between promoters and enhancers across liver and heart tissues, averaged over the four species. Percentages represent the fraction of overlapping elements in each category. **D.** Percentage of promoter and enhancer elements containing high-confidence orthologous regions across the four species (dark orange and blue bars). Lighter shade bars indicate promoters and enhancers with no high-confidence orthologs (non-alignable). Greylisted regulatory elements correspond to elements with elevated reads density in input ChIP (Methods). **E.** Sequence conservation of orthologous and non-alignable promoters and enhancers in mouse. Orthologous mouse elements have significantly higher phastCons sequence conservation scores than non-alignable elements. Both orthologous and non-alignable mouse elements have sequences significantly more conserved than the genomic background (Mann-Whitney U tests, with Benjamini-Hochberg correction for multiple testing, *** corrected p-values < 0.001). See also Figures S1, S2 and Table S1.

To compare regulatory regions across the four study species, we used pairwise wholegenome alignments to identify promoters and enhancers for which orthologous regions can be confidently assigned across all species (Figure 1A, and Methods). Specifically, we identified sub-regions of any promoter and enhancer with a strict alignment between the mouse genome and each of the three other rodents (Methods). To illustrate this process, we show regulatory elements in each species around the heart-specific genes *Myh6* and *Myh7*, for which a number of orthologous elements were identified (Figure 1A, solid boxes). Using this approach, we can compare regulatory activities at orthologous locations within a substantial fraction of promoters and enhancers across the four study species (shaded boxes and connecting lines in Figure 1A; Figure S2). Overall, and as expected for non-chromosomal genome assemblies, we identify 4-way orthologous regions within 20-30% promoters and enhancers in each species (Figure 1D). We define these elements as orthologous promoters and enhancers. In agreement with previous studies [22, 23, 26], orthologous promoters are mostly active across all study species (70%), whereas orthologous enhancers are largely active in only one species (55%, Fig. S2). Lastly, orthologous elements in either tissue show significantly higher levels of sequence conservation in mouse compared to non-alignable elements (Figure 1E). These results closely agree with those reported in recent similar datasets [33, 34], and indicate that across the four-species, we capture promoters and enhancers corresponding to both conserved and fast-evolving sequences. As such, the orthologous and non-alignable regulatory elements we defined constitute an exhaustive collection to study regulatory evolution in mole-rats.

### Phylogenetic modeling of regulatory activity identifies promoter and enhancer shifts in mole-rats

We next focused on orthologous promoters and enhancers to identify mole-rat specific changes in gene regulation. We adapted the EVE phylogenetic modeling method, initially developed for transcriptomic data [38], to identify promoters and enhancers showing shifts in regulatory activity along mole-rat branches (Figure 2A). This approach is based on an Ornstein-Uhlenbeck model with two optima, and allows statistical assessment of branch-specific shifts in regulatory activity while accounting for inter-species variation (Methods). Because EVE had not been applied to ChIP-seq data previously, we extensively evaluated the performance of phylogenetic modeling with simulated data (Figure S3 and Methods), and selected suitable data normalization and statistical thresholds (Figure S3 and Suppl. Table S2). Here, we focus on results with H3K27ac data, as it is broadly distributed across promoters and enhancers, and shows a large dynamic range of normalised read densities and good phylogenetic signal across our study species (Figure 2B and Figure S3).

**Figure 2:**
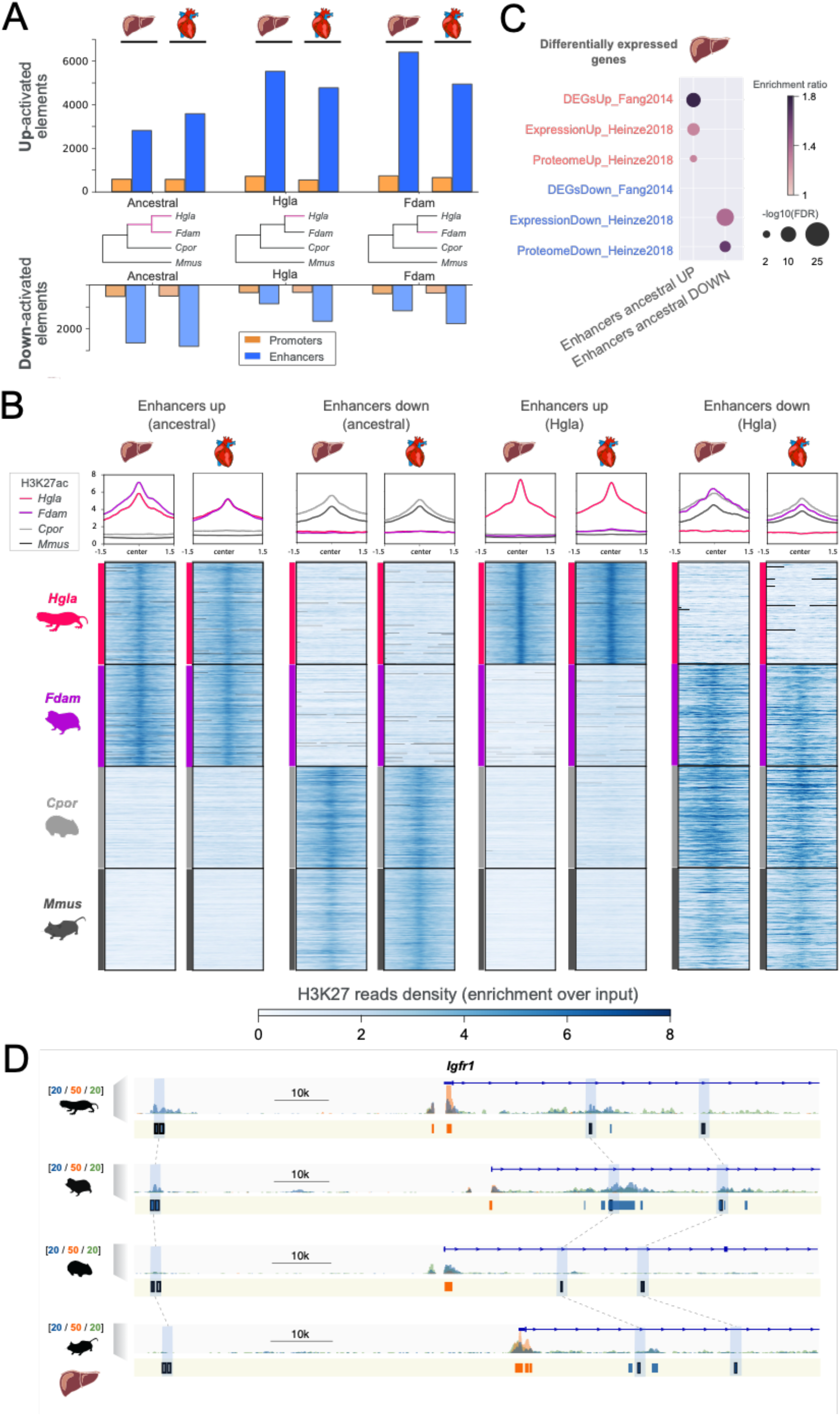
Identification of promoters and enhancers with an activity shift in mole-rats. **A.** Number of regulatory elements identified via phylogenetic modeling with an increase (Up) or decrease (Down) in activity in the indicated branches of the species tree: mole-rat ancestral (ancestral), naked mole-rat (Hgla) and Damaraland mole-rat (Fdam). **B.** H3K27ac reads density heatmaps and profile plots for up and down enhancers in the ancestral and naked mole-rat branches. Reads densities are presented as fold enrichment over input, averaged across biological replicates. **C.** Liver enhancers with an ancestral activity shift in mole-rats are significantly associated with previously identified differentially-expressed genes in mole-rats liver (GREAT enrichment tests, Methods). **D.** Example of liver enhancers up-regulated in the ancestral mole-rat branch and associated to the *Igf1r* insulin response locus. H3K27ac (blue), H3K4me3 (orange) and H3K4me1 (green) histone marks enrichment in liver around *Igf1r, scale units* as in Fig. 1A. Promoters are shown in orange, enhancers in blue. Up enhancers are identified by black boxes and linked by light blue boxes and dashed lines across species. See also Figures S3-S5, and Tables S2 and S3.

Using this approach, we identified shifts in regulatory activity across three branches of the study phylogeny (Figures 2A and B): the ancestral branch leading to mole-rats (Ancestral), and the single-species branches leading to the naked-mole rat (Hgla) and the Damaraland mole-rat branch (Fdam). Across both ancestral and single-species branches, phylogenetic modeling accurately identified orthologous regions presenting increased (“Up”) or decreased (“Down”) H3K27ac read densities in mole-rat species compared to outgroup rodents (Figure 2B). For each tested branch, phylogenetic modeling identified ~400 promoters and 3,000-6,000 enhancers with increased regulatory activity in mole-rat branches (Figure 2A, top bars) – which we define as Up elements. Conversely, we obtained ~200-500 promoters and ~1,000-2,500 enhancers with reduced regulatory activity (Figure 2B, bottom bars); which we define as Down promoters and enhancers.

We next compared the locations and properties of Up and Down elements identified with phylogenetic modeling (Table S3) to regulatory changes inferred by state-of-the-art methods, such as parsimony [33] and differential binding analysis [39] (Figures S4 and S5). Phylogenetic modeling recovered regulatory shifts with read density patterns clearly consistent with each tested branch (Figure 2B and Figures S4-5). Moreover, and to assess the relative significance of enhancer shifts inferred by the three methods, we overlapped their nearby genes with previously reported differentially expressed genes and proteins in mole-rat liver [13, 16] (Methods). In this analysis, enhancer shifts identified by phylogenetic modeling enriched more strongly in the vicinity of mole-rat differentially expressed genes (Figure 2C). In agreement with this observation, we found genes flanking Up and Down promoter and enhancer shifts include a number of previously reported *loci* with differential gene expression between mole-rats and other rodents, such as the upregulated insulin response gene *Igfr1* [13] (Figure 2D) or the downregulated uricase gene *Uox* [17] (Table S3).

### Enhancer shifts in the ancestral mole-rat branch enrich for tissue-specific metabolic and morphological adaptations

To investigate gene regulatory pathways associated with mole-rat enhancer shifts, we initially focused on the ancestral branch leading to both mole-rat species. On this branch, top enriched gene ontologies across enhancer shifts in liver and heart tissue provided a summary map of gene regulatory changes in mole-rats (Figure 3A and Tables S4-S6). In heart, Up enhancers were associated with heart contraction, energy metabolism and cellular respiration gene ontologies, consistent with the low heart rate and resting cardiac contractility observed in the naked mole-rat heart [40]. In liver, Up enhancers were linked to miRNA transcription, liver development and lipid metabolism, in line with reported adaptations in line with reported adaptations in fatty acid utilization in both mole-rat species [16, 41, 42]. For Down enhancers, top enrichments were observed for migration and angiogenic processes in heart, and hematopoietic and purine catabolism in liver – some of which align with previous reports [17]. In sum, while some of the associations observed above may reflect the biology of liver and heart tissues, several enrichments suggest specific connections to known adaptations in the mole-rat clade - and not observed in control analyses on the guinea pig branch (Table S7).

**Figure 3:**
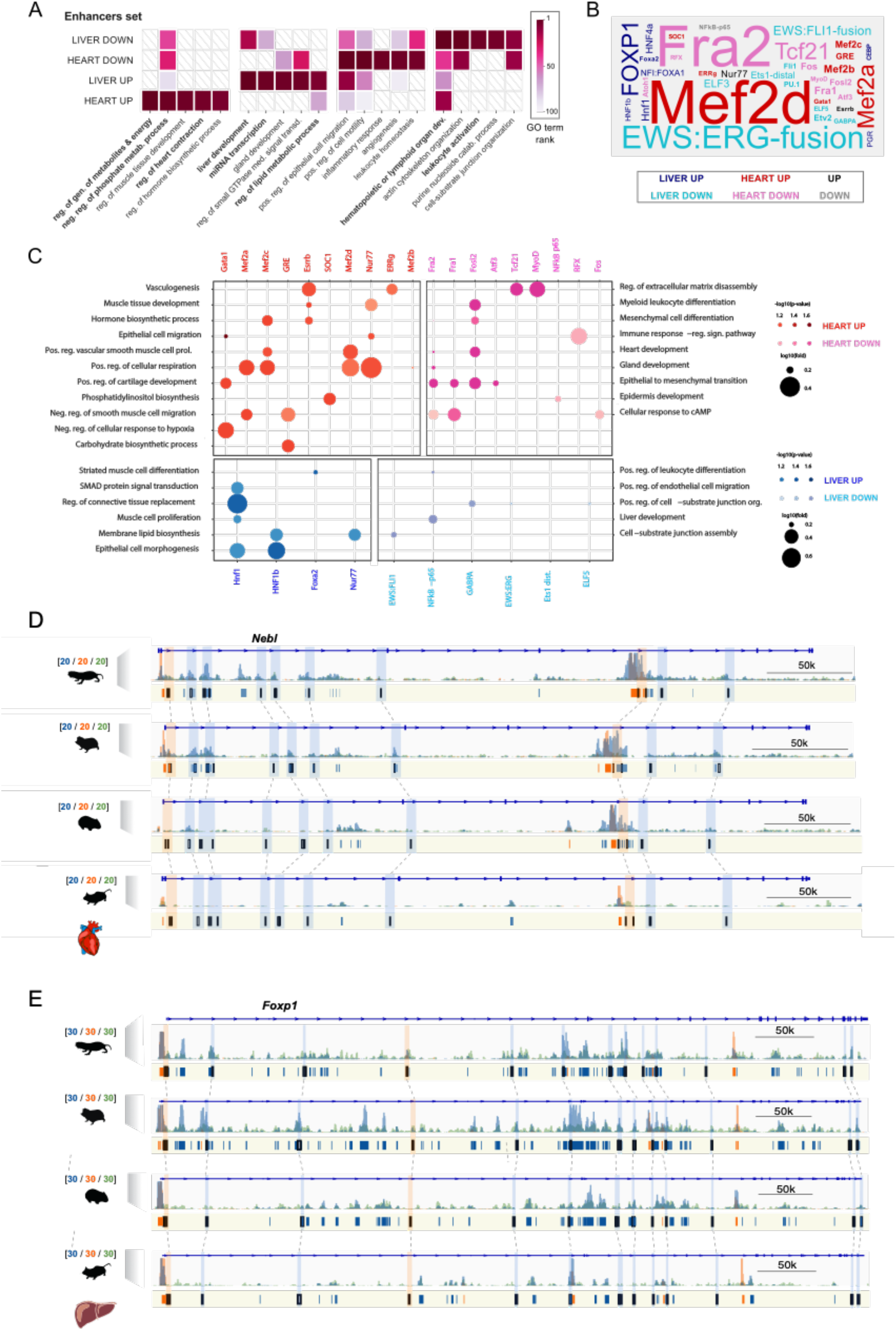
Gene ontologies and transcription factor binding sites enriched in enhancers with an activity shift on the ancestral mole-rat branch. **A.** Top gene ontology (GO) terms enriched in enhancers with an activity shift on the ancestral mole-rat branch. The top 5 associated gene ontology terms (GREAT enrichment tests, corrected p-values < 0.05, Methods) are shown for each set. Colors indicate the rank of the GO term in each set, while white barred boxes mark sets where the term is not enriched (not in the top 100). The complete list of enriched terms is available in Supp Tables S4-S6. **B.** Transcription factor binding sites (TFBS) enriched in mole-rat enhancers with an activity shift on the ancestral branch. Enriched TFBS are represented as a word-cloud, with colors indicating the enhancers set and size proportional to the rank of the TFBS in enrichment tests (HOMER enrichment tests, Methods). TFBS found enriched in up elements of both tissues are labeled ‘UP’, similarly for down. The complete list of enriched TFBS is available in Supp Table S7. **C.** Enriched transcription factor binding sites associated with specific gene ontology terms in heart and liver. Panels show significant associations between enriched TFBS and GO terms in shifted heart (top) and liver (bottom) enhancers, with up-activated enhancers on the left and down-activated enhancers on the right (hypergeometric tests, with Benjamini-Hochberg correction for multiple testing, corrected p-values < 0.1, Methods). GO terms for up enhancers are displayed on the left y-axis and GO for down enhancers on the right y-axis. **D.** Example of heart promoters and enhancers with an activity shift in mole rats and associated to the *Nebl* (Nebulette) locus. Representation as in Fig. 2D: H3K27ac (blue), H3K4me3 (orange) and H3K4me1 (green) histone marks enrichment in heart displayed around *Nebl*. Promoters are shown in orange, enhancers in blue. Orthologous up enhancers and promoters are identified by black boxes and linked by broken lines across species. **E.** Example of Up promoters and enhancers at the *Foxp1* locus in liver. Representation as in panel D. See also Tables S4-S9.

To further explore the relationship between enriched gene ontologies and tissue-specific gene regulation, we identified transcription factor binding sites (TFBSs) enriched in Up and Down enhancers in either tissue (Figure 3B and Table S8). Up enhancer shifts were enriched for TFBSs of tissue-specific transcription factors, such as MEF family members in the heart and FOX transcription factors in the liver. In contrast, Down enhancer shifts associated with TFBSs of broadly expressed transcription factors, such as FOS members in the heart and ETS proteins in the liver. Moreover, we detected specific pairs of TFBSs and ontology terms significantly associated with each set of enhancers (Figure 3C), suggesting regulatory rewiring of tissue-specific processes. As an example, for Up enhancers in the heart TFBSs for MEF family members associate with the cellular respiration ontology category, which includes genomic loci such as the coactivator *Ppargc1a*, a central regulator of mitochondrial biogenesis and respiratory capacity [43].

Taken together, these observations also suggest specific responses and transcriptional pathways with altered gene regulation in mole-rats (Table S9), such as the *Nebl* heart contraction locus (Figure 3D). In this region, we detect a large number of Up promoters and enhancers, some of which also contain MEF transcription factor binding sites. In liver, FOXP1 binding sites were enriched in Up enhancer shifts (Figure 3B), and we also found a concordant recruitment of Up promoters and enhancers at the *Foxp1* gene locus (Figure 3E, Tables S3 and S9), suggesting a coordinated upregulation of this pathway in mole-rats. FOXP1 is a known regulator of hepatic glucose homeostasis [44], and its regulatory rewiring in mole-rats may contribute to their enhanced glucose utilization compared to other rodents [7, 16].

### Enhancer shifts along mole-rat evolution identify temporal rewiring of gene regulation

Our study phylogeny allows for temporal investigation of regulatory evolution in mole-rats. We thus asked whether ancestral and species-specific enhancer shifts associate with common or divergent molecular processes along the evolution of mole-rats. To this end, we clustered related gene ontologies detected in enhancer shifts across branches (Methods; Figure 4A and Table S10), and we focused on gene regulatory pathways found across several branches in the phylogeny (Figure 4A).

**Figure 4:**
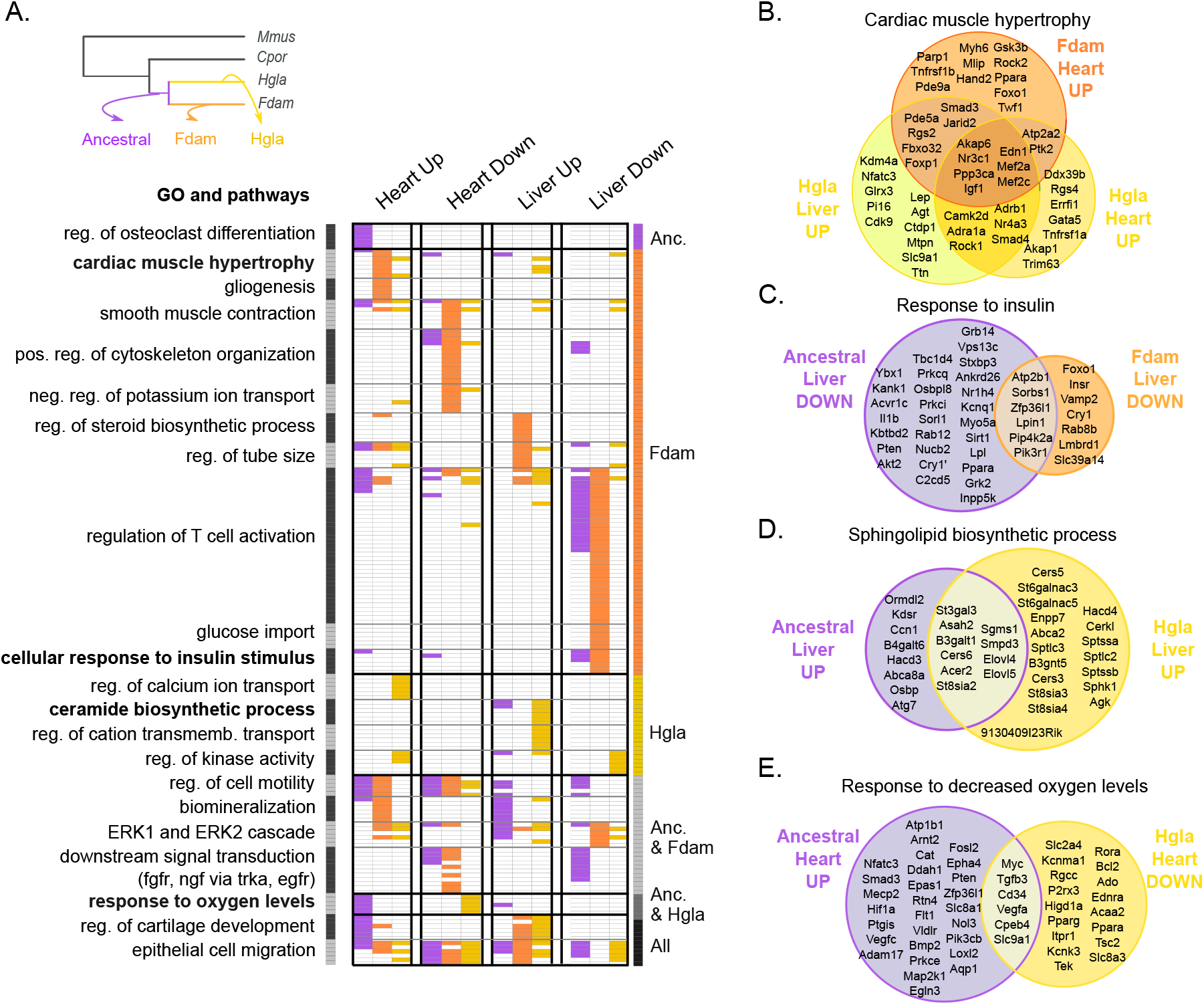
Comparison of gene ontologies and pathways enriched in enhancers with an activity shift in ancestral and single-species mole-rat branches. **A.** Gene ontology terms significantly associated to enhancers with activity shifts in mole rats across different branches (grey inset) and tissues. Similar gene ontology terms and pathways were grouped in clusters (left axis, Methods). Each row in the heatmap is a gene ontology term or pathway within a cluster, with its presence (colored) or absence (white) indicated in each enhancer set. Columns correspond to the different tissues and shift directions, and colors to the branches (grey inset). Clusters are ordered to highlight terms enriched in particular branches (right axis). The complete list of enriched terms and cluster membership are available in Supp Table S10. **B-E.** Selected examples of gene ontology terms enriched across enhancers of different branches or tissues. Venn diagrams show the overlap between genes associated with shifted enhancers of each set (Methods). See also Figure S6 and Table S10.

First, this comparison revealed temporal associations of gene ontologies and pathways along the three branches. We observe most gene ontologies enrichments were driven by one branch, such as cardiac muscle hypertrophy or response to insulin in the Damaraland molerat branch, and ceramide biosynthesis in the naked mole-rat branch. However, several processes appear to harbour regulatory rewiring on both ancestral and single-species branches, possibly in response to continued evolutionary pressure. We further explored some of these pathways by analysing whether enhancer shifts across branches are proximal to the same genes (Figure 4B-E).

Genes associated with cardiac muscle hypertrophy accumulate Up enhancers in both single-species branches, in the heart for Damaraland mole-rat and both in heart and liver for naked mole-rat (Figure 4B). Cardiomyocyte hypertrophy is a complex process mediated by signals arising from multiple cell types [45], including in the vasculature, the heart and the liver. Indeed, we found individual hypertrophy-related genes in each branch/tissue often had corresponding tissue-specific expression levels (Figure S6), such as *Myh6, Edn1* or *Trim63* in the heart, or *Agt* and *Igf1* in the liver. Thus our results suggest mole-rats have recently and convergently modified their response to cardiac hypertrophy, primarily in the heart but also at liver loci (such as the angiotensinogen locus *Agt* in the naked mole-rat branch).

Genes in the insulin response pathway are known to be altered in both the naked and the Damaraland mole-rats [13], with individual genes in this response being either induced or repressed. Down enhancers are the predominant gene regulatory change in this response, both in the ancestral and Damaraland mole-rat branches (Figure 4C). Although we detect a lower number of associated loci in the single-species branch, these include both upstream (*Insr*) and downstream (*Foxo1*) response genes, suggesting specific regulatory changes in this response in Damaraland mole-rat. Similarly, we found loci associated to ceramide biosynthesis in Up liver enhancers for both the ancestral and naked mole-rat branches (Figure 4D), which is consistent with reported high levels of short-chain ceramides in naked mole-rat brain lipids [46].

Lastly, we detected genomic loci associated with the response to hypoxia among heart enhancer shifts both in ancestral and naked mole-rat branches (Figure 4A and 4E). In agreement with the high protein levels of HIF1A in mole-rats [12], we detected Up enhancers in the ancestral mole-rat branch for both core regulators of hypoxic gene expression (*Hif1a, Epas1* and *Arnt2*) and *bona fide* HIF target genes (*Vegfa, Egln3, Aqp1*). However, we also found a number of hypoxia response gene loci harbour Down enhancers in the naked molerat branch. These results suggest that, in addition of an upregulation in the response to hypoxia in mole-rats, the naked mole-rat may also tune down gene regulatory landscapes for a subset of hypoxia-inducible genes. In support of this hypothesis, we tested whether regulatory changes in ancestral and mole-rat branches affect distinct sets of genes in the hypoxia response pathway (Figure 4E) and confirmed that loci associated with enhancer shifts in each branch overlap more rarely than expected at random (permutation test with 100,000 random resampling iterations, p-value = 0.009).

### Non-alignable mole-rat enhancers associate with SINE repetitive elements and rewire metabolic transcription factor networks

The phylogenetic modeling approach we developed here focuses on alignable segments within regulatory regions across our four study species. Nevertheless, we noticed some of the genomic loci previously linked to mole-rat traits are flanked by regulatory elements that cannot be aligned to the mouse genome (Figure S7), which we term non-alignable. Repetitive genomic regions (which are difficult to align across species) have been previously linked to rewiring of regulatory networks [33–35]. We thus investigated the genomic properties of non-alignable promoters and enhancers in our study, and their potential contribution to mole-rat specific gene regulation.

As expected, we observed non-alignable regulatory elements in mole-rats consistently enrich for repetitive element sequences estimated to be of young age (Figures 5A and B). Moreover, non-alignable enhancers significantly overlap with several repetitive element families (Figure S7). Among these, we identified enriched transcription factor binding sites suggestive of regulatory co-option (Figure 5C and Table S11). First, we found a consistent enrichment of RAR binding sites across non-alignable liver enhancers in all four species, and in sequences overlapping SINE/Alu repeats – which suggests rodent SINE/Alu elements contribute to RAR-mediated gene regulation, as previously proposed in primates [47]. In contrast, enrichment of HNF4A binding sites in SINE/ID repeats was specific to mole-rat liver enhancers in both species. HNF4-SINE/ID sequences enrich for monocarboxylic and fatty acid catabolism gene ontologies (Figure 5D), which include transcription factor loci with key roles in lipid metabolism, such as PPAR genes (Figure 5E). For RAR-SINE/Alu sequences enriched in non-alignable liver enhancers, we clustered gene ontology terms associated to their flanking genes across the four species (Figure 5F). In agreement with the known roles of RAR/RXR in fatty acid oxidation and lipid metabolism [48], we found terms related to lipid biosynthesis and catabolism consistently across all species. However, for mole-rats RAR-SINE/Alu sequences also associated with TGF-beta signaling and proteolysis, suggesting mole-rat specific repeats may rewire protein degradation and immunomodulation via the RAR network.

**Figure 5:**
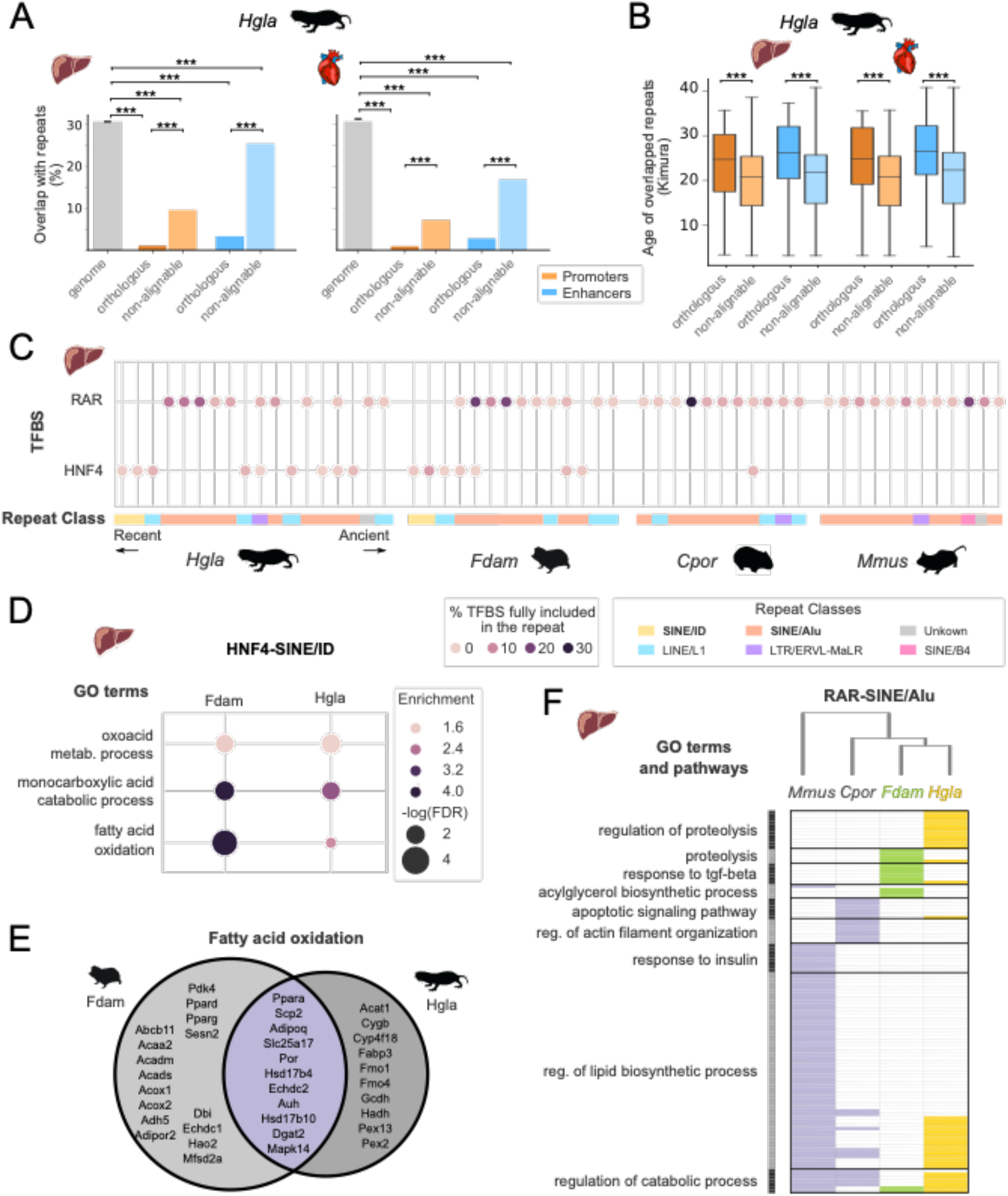
Non-alignable mole-rat liver enhancers associate with specific repetitive elements and transcription factor binding sites. **A.** Overlap with repetitive elements for naked mole-rat orthologous and non-alignable promoters and enhancers. Non-alignable enhancers and promoters overlap significantly more with repeats than orthologous elements, while all regulatory elements sets are significantly depleted in repeats compared to the genomic background (*** Benjamini-Hochberg corrected permutation-based p-values < 0.001, Methods). **B.** Non-alignable naked mole-rat promoters and enhancers overlap significantly younger repeats than orthologous promoters and enhancers (Mann-Whitney U tests, with Benjamini-Hochberg correction for multiple testing, *** corrected p-values < 0.001). Repeats ages were estimated with RepeatModeller (Methods), and are expressed as Kimura distances. **C.** Transcription factor binding sites (TFBS) for RAR and HNF4 in liver non-alignable enhancers significantly associate with specific repetitive elements. Circles indicate significant association between a specific repeat and a TFBS, with color intensity proportional to the fraction of total TFBS in non-alignable liver enhancers included in the repeat. Repeats are ordered by age and colored by repeat class. RAR TFBSs are significantly associated with SINE/Alu across all four species, whereas the HNF4 – ID SINE/ID enrichment is specific to mole-rats. **D**. Gene ontology terms significantly associated with non-alignable enhancers presenting a HNF4-SINE/ID instances in naked mole-rat and Damaraland mole-rat liver enhancers (GREAT enrichment tests, Methods). **E.** Venn diagram of genes associated to non-alignable mole-rat liver enhancers with HNF4-SINE/ID instances and annotated with the “fatty acid oxidation” gene ontology term. **F.** Gene ontology terms associated to RAR-SINE/Alu expansions across the four species. Representation as in Fig. 4A: gene ontology terms were grouped in clusters (shown on the left), absence (white) and presence (colored) of GO terms in each species are indicated. The complete list of enriched terms and cluster membership are available in Supp Table S10. See also Figure S7 and Table S11.

In conclusion, our results are consistent with a body of evidence on the importance of repetitive, transposon-derived elements providing a source of regulatory potential that can associate with functional innovation in mammals [33–35, 49].

## DISCUSSION

Evolutionary differences between mammals are expected to be largely driven by alterations in gene regulation rather than changes in protein sequences. Previous comparative studies of mammalian regulatory landscapes [23, 25, 26] have identified a continuum of regulatory regions from highly-conserved to lineage-specific [50], with the former being associated with pleiotropy and core tissue functions. However, the extent to which lineage-specific changes in regulatory activity contribute to tissue-level phenotypic adaptations is debated [51, 52], with the exception of a few well-characterised examples [35, 53–55].

To date, most comparative analyses of regulatory landscapes have been based on presence or absence of orthologous promoters and enhancers, and thus have a number of methodological limitations [36, 37]. These approaches do not normalise or quantitatively compare epigenomic levels of promoter and enhancer activity across species, and typically do not account for varying phylogenetic distances between species. To extend and improve these analyses, we applied and validated a phylogenetic modeling approach to promoter and enhancer epigenomic activities in two tissues from a four species phylogeny – in which we compare two mole-rat species with guinea pig and mouse as outgroup rodents. This strategy allowed us to quantitatively identify lineage-specific shifts in promoters and enhancers across mole-rat branches, investigate the associated biological processes, and assess the contribution of mole-rat specific changes in gene regulation to their characteristic evolutionary traits. The method we describe here improves on previously known issues with comparative analyses of functional genomics data, and could be extended for similar datasets across species, tissues and developmental stages. Our work adds to ongoing efforts to extend statistically-sound, quantitative evolutionary models to the analysis of functional genomics data [34, 36, 56, 57].

Our results reveal several contributions of mole-rat gene regulation to tissue-specific processes. First, we find a consistent upregulation of enhancers associated with *Ppar/Pgc1a* loci, especially in the heart, but also in the liver. These mole-rat specific changes are consistent with the role of PGC1a-PPARs in lipid metabolism and mitochondrial biogenesis. Several studies have documented alterations in mitochondrial function [16, 42] and morphology [58, 59] in mole-rats, which our results support from an epigenomic perspective. Moreover, we found lineage-specific liver enhancers in mole-rats enrich for SINE repeat families and associate with fatty acid oxidation processes, which is in agreement with the reported increase in fatty acid utilization in mole-rat liver [16]. Also in liver, we detected upregulated mole-rat enhancers in the *Foxp1* locus, as well as an overrepresentation of FOXP1 binding sites across upregulated liver enhancers. Our observations suggest mole-rat specific changes in the FOXP1 regulatory network, consistent with its known functions as a repressor of liver gluconeogenesis [44] and the reported high rates of glucose utilization in mole-rat tissues [16]. In heart, some of the strongest upregulated enhancers associate with myocardial conduction processes, and include genes such as *Nebl, Cacna1c and Myh7*. These changes in heart-specific gene regulation in mole-rats likely contribute to the morphological and conduction properties of the mole-rat myocardium [40, 60].

Comparing across ancestral, naked mole-rat and Damaraland mole-rat branches, our results identified recurrent regulatory changes associated to specific pathways. We found upregulated enhancers associated to cardiac hypertrophy both in the Damaraland mole-rat and naked mole-rat branch; mostly in the heart but also in the liver, and including mole-rat specific enhancer gains in loci such as *Igf1, Agt* and *Foxo1*. This observation suggests a complex landscape of mole-rat changes in this pathway that may inform reduced levels of hypertrophy in mole-rat myocardium [10, 60]. In contrast, for enhancers associated with the insulin response, we found a consistent downregulation in mole-rat liver, both in the ancestral and Damaraland mole-rat branches. Previous work documented both upregulated and downregulated expression of insulin response genes in mole-rats [13], and our findings suggest epigenomic enhancer downregulation is particularly significant in this response. Lastly, for loci involved in the response to hypoxia, we found evidence of both epigenomic upregulation in the ancestral branch and downregulation in the naked mole-rat branch. The former is consistent with the known changes in HIF1A protein sequence in mole-rats resulting in reduced proteasomal degradation [12]. Additionally, the epigenomic enhancer downregulation we observe in the naked mole-rat branch suggests a second wave of rewiring in this response that may fine-tune hypoxic regulation at a subset of loci. Although speculative, a possible interpretation is that downregulated naked mole-rat enhancers in this response may enrich for HIF target genes, as suggested by their overlap with hypoxia-inducible genes in humans (70%; [61]) and lower affinity in HIF binding sites compared to mouse (Figure S6). Nevertheless, further experimental work would be required to substantiate this hypothesis.

There are a number of limitations to our approach. First, our ability to compare epigenomic signals across species is partly dependent on reference genome assemblies, alignments and annotations, which are of variable quality and completeness across our study phylogeny.

Second, we identified promoter and enhancer shifts based on epigenomic enrichment of H3K27ac, which strongly associates with canonical promoter and enhancer elements across the genome. H3K27ac levels display strong phylogenetic signal across species, thus representing a good quantitative proxy of regulatory evolution. Moreover, enhancers harbouring this histone mark have been experimentally validated in developmental models, and over 60% drive expected expression patterns [22, 62]. Nevertheless, non-canonical enhancers are increasingly recognized as an additional source of regulatory potential [63]. Such enhancers are often not associated with H3K27ac levels and thus invisible to our approach, potentially limiting the completeness of the lineage-specific shifts we identified here. Third, our phylogenetic modeling approach is affected by the size of the study phylogeny, which is constrained by available genomic resources in this clade, and sample accessibility limitations for wild mole-rat species. Thus our data represents a strategic compromise between phylogenetic coverage and epigenomic completeness. Nevertheless, it is likely a denser or deeper phylogenetic sampling would impact performance of this approach, by improving sensitivity and reducing false discovery estimates. Similarly, our analyses of non-alignable promoters and enhancers in mole-rats rely on our ability to identify lineage-specific sequences in the study phylogeny, which would be enhanced by a denser phylogenetic sampling in this clade.

In summary, this study presents a quantitative phylogenetic framework with which to investigate the long-standing question of how lineage-specific changes in gene regulation can contribute to phenotypic traits. By modeling epigenomic activities of promoters and enhancers within the mole-rat clade, we demonstrate the utility of this approach to identify promoter and enhancer shifts across ancestral and single-species branches, and connect these lineagespecific innovations with candidate tissue-specific processes rewired in mole-rats.

## METHODS

### Chromatin immunoprecipitation and high-throughput sequencing

We performed chromatin immunoprecipitation experiments followed by high throughput sequencing (ChIP-seq) using liver and heart tissue samples isolated from naked mole-rat, Damaraland mole-rat, guinea pig and mouse. The origin, number of replicates, sex, and age for each species’ samples are detailed in Table S1.

All animal experiments conducted in the UK were in accordance with the UK Animals (Scientific Procedures) Act 1986 Amendment Regulations 2012 and performed under the terms and conditions of the UK Home Office licenses P51E67724 (D.V.) and P7EBFC1B1 (E.J.S.). Damaraland mole-rat tissues were obtained from the Kuruman River Reserve, Kalahari Research Trust following local ethical approval (export permit FAUNA 0718/2/2016) and imported to the UK (DEFRA import authorization ITIMP16/0374).

At least two independent biological replicates from different animals were performed for each species and antibody. Wherever possible, tissues from young adult males were used. Tissues were prepared immediately post-mortem (typically within an hour) to maximize experimental quality. Post-mortem tissues were kept on ice until processed to minimize potential loss of protein-DNA interactions during post-mortem time. Sample allocations to experimental batches were randomised to ensure unbiased distributions of species, tissue, individual and sex, using the R/Bioconductor package OSAT [64].

Tissues were prepared by direct perfusion of the liver with PBS, followed by dicing the whole organs (liver and heart) in small pieces around 1cm^3^. Blood clots within the heart ventricles were removed. Cross-linking of the diced tissue was performed in 1% formaldehyde solution for 20 min, addition of 250 mM glycine and incubation for a further 10 min to neutralize the formaldehyde. After homogenization of cross-linked tissues in a dounce tissue grinder, samples were washed twice with PBS and lysed according to published protocols [65] to solubilize DNA-protein complexes. Chromatin was fragmented to 300 bp average size by sonication on a Misonix sonicator 3000 with a 418 tip (1/16 inch diameter). Chromatin from 50-200 mg of dounced tissue was used for each ChIP experiment using antibodies against H3K4me3 (millipore 05-1339), H3K27ac (abcam ab4729) and H3K4me1 (abcam ab8895). Illumina sequencing libraries were prepared from ChIP-enriched DNA using ThruPLEX DNA-seq library preparation kit (Takara Bio) with up to 10ng of input DNA and 8-15 PCR cycles. After PCR, libraries were pooled in equimolar concentrations and sequenced on Illumina HiSeq 4000 or NovaSeq instruments.

### Computational analysis of ChIP-seq data and definition of regulatory regions

#### Basic alignment and peak calling

Aligned bam files were obtained with bwa 0.7.17 and the Ensembl v99 assemblies HetGla_1.0 (Naked mole-rat), DMR_v1.0 (Damaraland mole-rat), Cavpor3.0 (guinea pig) and GRCm38 (mouse). We used macs2 [66] to call peaks for each ChiP-Seq replicate, using default parameters and “--keep-dup all” to retain duplicate reads. Before peak-calling, multi-mapping reads were removed and read-depth adjusted to 20 million uniquely mapped reads (or all available reads for low-depth libraries).

#### Definition of regulatory regions from ChIP-seq peaks

We first constructed sets of reproducible peaks for each combination of histone mark, tissue and species by merging peaks identified across a minimum of two biological replicates (with minimum 50% length overlap). In a second step, we used these sets of reproducible peaks to identify promoters, enhancers and primed enhancers independently for each species and tissue, according to the following criteria: all H3K4me3 peaks overlapping an H3K27ac peak (minimum 50% bases overlap) were predicted to be promoters, H3K27ac peaks not overlapping promoters were predicted as enhancers, and H3K4me1 peaks not overlapping any H3K4me3 or H3K27ac peaks were as predicted primed enhancers.

### Cross-mapping of regulatory regions across four rodent species

#### Definition of orthologous regulatory regions

we used the LiftOver local software [67] to map genomic coordinates of the regulatory elements identified in naked mole-rat, Damaraland mole-rat and guinea pig to the mouse genome. LastZ pairwise whole-genome alignments with mouse were downloaded from Ensembl Compara v99 and converted from ensembl maf format to the UCSC chain format using UCSC tools (including mafToPsl, pslToChain and chainSwap, v357 for all software). We validated the correct implementation of the coordinates conversion step using LiftOver with the generated chain alignment files by comparing the resulting coordinates with those obtained from queries directly performed through the Ensembl API[68]. We defined a set of “high-confidence orthologous regions” as 4-way orthologous regions for which we required robust LiftOver mapping of regulatory elements across the four genomes. This was achieved in two steps, involving (i) the definition of the set of regulatory regions, expressed in mouse genome coordinates, that can be aligned from each of the other species to mouse; and (ii) a filtering step to retain only (sub)regions with a strict reciprocal LiftOver mapping in each genome (e.g. a naked mole-rat region mapping to mouse, and the mouse coordinates of this region mapping back to each of the other three genomes). In this filtering step, the final criteria for regions to be defined as high-confidence 4-way orthologs were: a reciprocal LiftOver mapping from mouse to the three other genomes, with a minimum of 30% coverage (-minMatch 0.30) and similar lengths of the resulting regions across the four genomes (the difference in length between the mouse region and regions in any other species must be less than 15% of the largest region).

#### Homogenization of regulatory region type

for cases where orthologous regions were defined as different types of regulatory elements across species, we used a majority rule system to homogenize regulatory element types. For instance, a region defined as a promoter in mouse, guinea pig and naked mole-rat but as an enhancer in Damaraland mole-rat was re-defined as a promoter across all species. In case of ties, regions are arbitrarily assigned to the “highest-level” regulatory type (i.e. promoters have priority over enhancers and primed enhancers, and enhancers have priority over primed enhancers).

#### Exclusion of greylisted regions

H3K27ac signal read density normalization with input ChIP is critical for quantitative analyses of regulatory activity. We therefore established greylists of regions with an unusually elevated signal in input ChIP experiments to tag them for exclusion. We took advantage of the approach implemented in the GreyListChIP R package, which flags regions with elevated signal in the input. In practice, for all pairs of input – H3K27ac sample ChIP experiments, we ran chipseq-greylist v1.0.2 with 100 bootstrap (--bootstraps 100), a simpler python implementation of GreyListChIP (available from https://github.com/roryk/chipseq-greylist). Two greylists were computed, one for each tissue, and including all regions flagged by chipseq-greylist in at least one input ChIP in any species. Finally, any region from the set of “high-confidence orthologous elements” overlapping with a greylisted region was removed before phylogenetic modeling. Across the different sets, the fraction of alignable elements that we exclude as greylisted ranged between 0.5% and 2%.

### Sequence conservation of orthologous and non-alignable regulatory elements

We downloaded sequence conservation scores (“phastCons”) for the mouse genome from the UCSC (http://hgdownload.cse.ucsc.edu/goldenpath/mm10/phastCons60way/), which were computed from a multiple alignment of 60 vertebrate genomes. We used the UCSC tool bigWigAverageOverBed (version 357) to extract average phastCons scores over each regulatory region included in the “non-alignable”, “orthologous”, and “genome” regulatory element sets. The “Random” element sets were built from 100 permutations of the “non-alignable” element sets on the mouse genome, excluding exons.

### Phylogenetic modeling of shifts in regulatory activity on mole-rat branches

#### Reads density normalization for phylogenetic modeling

we first normalize the H3K27ac FPKM signal for each library with the corresponding input control, retaining the log2 fold change of signal FPKM over input FPKM. Specifically, we extracted H3K27ac reads density (FPKM) at orthologous enhancers and promoters for each replicate and its corresponding input control experiment with the BAMscale cov utility [69], retaining only confidently mapped reads (-q 13). We next used quantile normalization to normalize fold changes across species and replicates, and verified that samples group according to the species phylogeny after normalization (Figure S3). This normalized data serves as the basic input for phylogenetic modeling of regulatory activity across species.

#### Phylogenetic modeling and detection of regulatory activity shifts

We modelled the evolution of regulatory activity along the study phylogeny using the EVE model, an improved Ornstein-Ulhenbeck model for continuous trait evolution under selection and drift [38]. Specifically, the EVE’s null model has four parameters that govern how traits evolve: the σ parameter (strength of drift), the α parameter (strength of selection), the θ parameter (optimal trait value) and the β parameter (ratio of intra- to interspecies variation). To detect significant shift in regulatory activity of enhancers and promoters on specific branches of the study phylogeny, we took advantage of EVE’s branch shift test. The branch shift test conducts Likelihood Ratio Tests (LRT) comparing two nested models: the four-parameters null model to a five-parameters model with a branch-specific shift in the optimal trait value. In the branch model, foreground branches have a distinct optimum (θ_1_) from background branches (θ_0_). By leveraging replicates and explicitly accounting for within-species variation (with the parameter β), the EVE model significantly improves upon classical OU models, as it reduces false positive inferences of stabilizing selection and optimum shifts [37, 38]. A challenge to applying EVE’s branch shift test to ChIP-seq data is its relatively low statistical power, typically requiring a large amount of data (number of replicates and size of the phylogeny) that is unusual even in RNA-seq experiments [70]. In addition, on small phylogenies, values of the test statistic (Likelihood Ratio Tests, LRT) can depart from the chi-square distribution [38]. Therefore, we performed computational simulations to (i) determine the distribution of LRT in our data and compute accurate p-values; and (ii) evaluate the true positive rate and select an appropriate alpha threshold for the branch shift test.

We ran EVE’s branch shift test [38] over normalized ChIP-seq reads using the evemodel R package [71] to detect regulatory activity shifts on the following branches of the four-species phylogeny: the ancestral mole-rat branch (Ancestral), the naked-mole rat branch (Hgla) and the Damaraland mole-rat branch (Fdam). Branch lengths in substitutions per site were extracted from Ensembl Compara v99 [68] (specifically, from the species tree computed from pairwise whole-genome alignments, available at https://github.com/Ensembl/ensembl-compara/blob/release/99/conf/vertebrates/species_tree.branch_len.nw). For each of the four orthologous regulatory elements sets (Enhancers Heart, Enhancers Liver, Promoters Heart and Promoters Liver), we first estimated model parameters under the null model (selection without shift), in order to obtain realistic parameter values for simulations. We then performed n=1,000 simulations under the null model for each of the four sets, using the mean value of the estimated model parameters (alpha, beta, theta0 and sigma). We next tested whether the study phylogeny was sufficiently large for Likelikood Test Ratios (LRT) to be distributed as a chi-square distribution: we computed LRT comparing likelihoods of the simulated data under the null model and under a model with a shift in the ancestral mole-rats branch, and found that LRT did not follow a chi-square distribution. We thus used these simulations under the null model to compute empirical p-values (Figure S3), as recommended previously[38].

Because the branch shift test has relatively low statistical power, we performed additional simulations to evaluate the false positive and true positive rates under a variety of simulation settings, and to select a suitable alpha threshold for our test statistic. We again performed n=1,000 simulations for each of the four sets (Enhancers Heart, Enhancers Liver, Promoters Heart and Promoters Liver), and with varying proportions of simulations under the null and shift models (from 5 to 17.5% of simulations with a shift). We performed simulations under the null and shift models using the average parameter values estimated from the data (parameters alpha, beta, theta0 and sigma), while the additional shift parameter value for the simulations with shift were drawn from a beta distribution Beta(alpha=8, beta=2)*3, selected to resemble empirically-estimated shift values. On average and across the four sets, testing for shift on the Ancestral branch at alpha = 0.05 and without filtering on the value of the estimated shift yielded a true positive rate of 0.46. We found that selecting alpha = 0.20 and filtering on absolute shift values > 1.5 yielded an acceptable false positive rate (0.08), while increasing the true positive rate to 0.77 (Table S2). Similar results were obtained with the corresponding simulations for shifts in each of the naked mole-rat (Hgla) and Damaraland mole-rat (Fdam) branches (Table S2).

Based on the above, for the four orthologous regulatory elements sets (Enhancers Heart, Enhancers Liver, Promoters Heart and Promoters Liver) and each of the three tested branches, we obtained p-values for the branch shift test and estimated shift values for each regulatory region. We then retained as differentially active elements all elements with empirical p-value < 0.2 and absolute shift value > 1.5. Reads density heatmaps and profile plots were drawn with Deeptools version 3.5.0 [72].

### Detection of regulatory activity shifts using parsimony

For comparison with phylogenetic modeling, we identified up and down promoters and enhancers in mole-rats from the same set of high-confidence orthologous elements with a complementary parsimony-based approach. To achieve this, we applied the dollo parsimony criterion, which states that a trait – here the regulatory activity of a genomic region - can be gained only once but can be lost multiple times. Under this criterion, and using the ancestral mole-rat branch as an example, Up enhancers are the enhancers detected as active in both mole-rats (i.e. overlapping a MACS2 H3K27ac peak corresponding to an enhancer in both species) and inactive in both guinea pig and mouse. Respectively, Down elements are detected as inactive in both mole-rats and as active in both guinea pig and mouse. Therefore, and in contrast to phylogenetic modeling, parsimony-based regulatory shifts are based on binary (presence or absence of peaks) rather than continuous data (normalised reads).

### Detection of regulatory activity shifts using differential binding analysis

For comparison with phylogenetic modeling, we implemented the procedure described in the DiffBind R package [39] to identify regulatory regions differentially active in mole-rats compared to guinea pig and mouse. For each regulatory region, we measured regulatory activity as the raw number of H3K27ac ChIP-seq reads minus the raw number of input ChIP-seq reads. We then normalized these counts across samples using the median of ratio method from the DESeq2 package [73]. Finally, we ran the DESeq2 differential analysis procedure, contrasting a first group containing all samples from both mole-rats against a second group with samples from guinea pig and mouse. We filtered DESeq2 results to retain differentially active elements with FDR < 0.01 and abs(log2foldchange) > 2.

### Gene ontologies, pathways and genes enrichment tests

We downloaded the mouse gene ontologies (“GO”) data from the MGI database on the 20^th^ of April 2022, as well as C2 pathway annotations from MgDB on https://bioinf.wehi.edu.au/software/MSigDB/. We filtered GO data to retain only GO of the Biological Process (“BP”) domain, and C2 pathways to retain only 1,397 pathways (mostly retaining REACTOME, KEGG and BIOCARTA pathways). We transferred mouse gene ontology and pathway annotations to the naked mole-rat and Damaraland mole-rat using gene orthologies from ensembl compara version 102, extracted with ensembl biomart [68]. We re-implemented the region-based gene ontologies tests implemented in GREAT [74], allowing us to use custom genomes (the naked-mole rat and Damaraland mole-rat genomes) and the more recent gene ontology annotations downloaded from MGI.

We used GREAT’s default gene to regions association rule, establishing gene basal regulatory domains (5kb upstream, 1kp downstream) with distal extension up to 1kb (or to the nearest next basal domain if it is closer). GREAT can perform three GO enrichment tests: a binomial test over regions against the genome as background (a), a hypergeometric test over genes against the genome as background (b) and a hypergeometric test over regions using a carefully selected set of genomic regions as background (c). We implemented all three tests, using similar rules for GO propagation and filters as in GREAT. We propagate GO annotations by associating genes annotated to a specific GO to all of this GO term’s parents. To increase statistical power and alleviate redundancy, we filter GO to only test for the most specific GO terms amongst all GO associated to the exact same foreground genes. For gene ontology and pathways enrichment tests, we used GREAT recommended tests (a) and (b) (i.e. tests against the whole genome as background). We retained only terms found enriched with both tests, ranked them according to BH adjusted p-value < 0.05 of test (a), and selected the top 100 GO terms and top 20 C2 pathways. To identify single instances of significantly-enriched genes we used test (c), using the complete regulatory regions set of a given category as background (for instance all orthologous heart enhancers as background for nmrdmr up heart enhancers). Lastly, we validated our implementation by verifying that highly similar terms were found when using the GREAT web-server with the mouse genome and older gene ontology data from MGI. For regulatory elements with an activity shift in the ancestral mole-rat branch, we used the Damaraland mole-rat genome, which has a higher contiguity, to perform functional enrichment tests. For single-species shifts and elements enriched in repeats, we used this species genome for the tests.

### Transcription-factor binding sites (TFBS) enrichment analysis

We used the HOMER software suite version 4.11 [75] to identify enriched TFBS in sets of regulatory regions (findMotifsGenome.pl script with default parameters, with “-h” to conduct hypergeometric tests and “-size given” to only search for motifs within regulatory regions). We used the Damaraland mole-rat genome as background to search for enriched TFBS in the regulatory elements with an activity shift on the ancestral branch. We retain the TOP 10 enriched TF for each element set (all BH-corrected p-values <0.05).

We conducted hypergeometric tests to identify significant TFBS-GO term associations amongst the top 10 TFs and top 100 GO enriched in each element set. For each GO-TFBS pair, we counted (i) the number of regulatory elements both containing the TFBS and associated with the GO, and (ii) computed the enrichment ratio compared to the expected number of overlaps. To alleviate the burden of multiple testing, we performed the hypergeometric test only for ratios > 1.5 and then retained all TFBS-GO pairs with a BH-corrected p-value < 0.1.

### Clustering of GO terms for visualization of enrichment across branches and tissues

To compare functional enrichments across sets of shifted regulatory elements in different tissues and branches of the phylogeny, we first removed redundancies by filtering out from each set the elements found shifted in two branches (i.e. “ambiguous” elements, found shifted in the ancestral and in one mole rat branch). We next used the GREAT approach to identify enriched GO terms and pathways in each filtered element set, retaining the top 100 GO (all corrected p-values < 0.05) and 20 C2 (all corrected p-values < 0.1) pathways (see “Gene ontologies, pathways and genes enrichment tests” for implementation details). For visualization, we grouped together similar GO terms and pathways found within or across sets, based on the overlap of associated genes. Specifically, two GO (pathways) were grouped in the same cluster if the jaccard index of the overlap was > 0.35. We next relax the jaccard index threshold to 0.2, to tentatively connect remaining singletons GO to existing clusters. We retained all clusters of size > 3 GO terms (and/or pathways) for visualization (Figure 4A).

### De novo repeat annotation in mole-rat species

We constructed *de novo* repeat libraries for naked mole-rat and Damaraland mole-rat using RepeatModeler version 2.0.2a [76] with default parameters. For guinea pig and mouse, we downloaded pre-computed *de novo* repeat libraries from the Dfam database [77]. We next ran RepeatMasker version 4.1.2-p1 to annotate the location of repeats in mole-rat, guinea pig and mouse genomes, in sensitive mode (-s) and skipping bacterial insertion check (-no_is), with rmblastn version 2.10.0+. The average 2-parameters Kimura distance of TE sequences to their consensus were computed using the calcDivergenceFromAlign.pl script from RepeatMasker, which corrects for elevated mutation rates in CpG loci. Finally, landscape plots were drawn using the createRepeatLandscape.pl script from RepeatMasker.

### Overlaps between regulatory elements and repeats

We used bedtools version v2.29.2 [78], across “orthologous”, “non-alignable” and “genome” elements sets. Specifically, we used bedtools intersect to compute overlaps between elements and repeats; and bedtools jaccard to obtain the corresponding jaccard index (ratio of the intersection to the union of the datasets, in number of bases). “Genome” sets were constructed from n=10,000 random permutations of the corresponding “orthologous” or “non-alignable” set, on the whole genome. Following the approach from the jaccard test of the GenomTriCorr R package [79], we tested for significant differences in overlaps with repeats across sets. Here, for comparisons with “genome” elements, the n=10,000 permutation-based jaccard indexes form the null distribution to which we compare the jaccard obtained on the corresponding “orthologous” (or “non-alignable”) set. For comparisons between “orthologous” and “non-alignable” sets, we compare the jaccard obtained on a “non-alignable” set with n=m elements to the null distribution obtained from 100 random samplings of n=m elements drawn from the “orthologous” and “non-alignable” sets combined. P-values were adjusted for multiple testing using the Benjamini-Hochberg (BH) procedure.

We again used the jaccard test to define significantly enriched repeat families. To do this, we first selected repeat families with a minimum of 100 instances overlapping an element in a “non-alignable” set. Second, for each repeat family, we computed the jaccard index of the intersection between regulatory elements of a “non-alignable” set and the repeats in a family. Third, we constructed a null distribution of n=100 jaccard indexes computed from the random permutations of “non-alignable” elements over the genome. Finally, we retained repeat families as significantly enriched in a “non-alignable” set when BH adjusted p-values were < 0.05.

### Enrichment of TFBS in repeats

We identified enriched TFBS in the set of non-alignable enhancers of each mole-rat using the HOMER software (see “Transcription-factor binding sites (TFBS) enrichment analysis”). We again used a permutation of intervals approach to identify TFBS significantly found fully included in a repeat more often than expected by chance. The null distribution was obtained from n=100 random permutations of TFBS within non-alignable enhancers. In mole-rats, we tested for significant association between enriched repeats and the TOP 10 enriched TFBS. In the outgroups, we tested only for significant association between HNF4 and RAR TFBS and enriched repeats.

Finally, we used GREAT (see “Gene ontologies, pathways and genes enrichment tests” for details) to test for functional enrichment for enhancers with the co-occurring RAR TFBS and Sine/Alu repeats and HNF4 TFBS and Sine/ID repeats.

## Code and data availability

The ChIP-sequencing data reported here has been deposited in GEO (accession number GSE222972).

All original code and datasets generated in this study have been deposited in Zenodo (https://doi.org/10.5281/zenodo.7442105). Briefly, these include all scripts, inputs and environments used to: (i) predict regulatory elements from histone mark peaks, (ii) identify orthologous elements across the 4 rodents, (iii) compute normalized H3k27ac read densities for orthologous elements (iv) identify shifted elements in mole-rats with phylogenetic modeling and (v) conduct gene ontology enrichment tests for mole-rats regulatory elements. All included supplemental datasets are listed in the Supplemental Material.

## AUTHOR CONTRIBUTIONS

E.P., D.V. and C.B. designed experiments; D.V., S.F. and A.U. performed experiments; E.P., D.V. and C.B. analysed the data; T.J.P., M.Z. and E.J.S. provided tissue samples; D.V., E.P. and C.B. wrote the manuscript. All authors read and approved the final manuscript.

## ACKNOWLEDGEMENTS

We thank the QMUL and CRUK-CI Genomics cores, David Gaynor and Katy Goddard (Kalahari Research Trust), and Nigel Bennet (University of Pretoria) for technical assistance; Pierre Vincens (IBENS, Paris), Bronwen L Aken (European Bioinformatics Institute) and Queen Mary’s Apocrita HPC facility, supported by QMUL Research-IT, for the coordination of computing resources; and Alexandre de Mendoza (QMUL) and Masa Roller (European Bioinformatics Institute) for critical comments on the manuscript.

