## Supplementary material for "Phylogenetic modeling of enhancer shifts in African mole-rats reveals regulatory changes associated with tissue-specific traits": Suplemental data

**SUPPLEMENTAL INFORMATION**

***Phylogenetic modeling of promoter and enhancer landscapes in mole-rats reveals gene regulation adaptations***
Parey et al. 2023

---

**SUPPLEMENTAL DATA**

**Figure S1. Basic properties of promoters, enhancers and primed enhancers identified in heart and liver tissues across two mole-rats and two outgroup rodents** (related to Figure 1) **(page 3)**

**Figure S2. Cross-mapping of promoters, enhancers and primed enhancers across the four study species** (related to Figure 1) **(p. 4)**

**Figure S3. Data normalisation and parameter estimation for phylogenetic modeling** (related to Figure 2) **(p. 5)**

**Figure S4. Comparison of phylogenetic modeling results in heart with regulatory shifts inferred from parsimony and differential binding analyses** (related to Figure 2) **(p. 7)**

**Figure S5. Comparison of phylogenetic modeling results in liver with regulatory shifts inferred from parsimony and differential binding analyses** (related to Figure 2) **(p. 8)**

**Figure S6. Additional properties of ancestral and single-species enhancer shifts in selected ontology categories** (related to Figure 4 and Discussion) **(p. 10)**

**Figure S7. Additional properties of repetitive elements in mole-rat genomes and their association with non-alignable enhancers** (related to Figure 5) **(p. 12)**

**Table S1**: Species and tissue samples used in this study (related to Methods, Figure 1 and Figure S1) **(p. 14)**

**Table S2.** Simulations to calibrate statistical thresholds for the branch-shift test of the phylogenetic modeling approach (related to Figure 2) **(p.14)**

**Table S3.** Genes significantly associated to enhancers with an activity shift in the ancestral mole-rat branch (related to Figure 2) **(p.14)**

**Table S4.** Gene ontology enrichments (TOP 100) for enhancers with an activity shift in the ancestral mole-rat branch (related to Figures 3 and 4) **(p.14)**

**Table S5.** C2 pathway enrichments for enhancers with an activity shift in the ancestral mole-rat branch (related to Figures 3 and 4) **(p.14)**

**Table S6.** Gene ontology enrichments for promoters with an activity shift in the ancestral mole-rat branch (related to Figures 3 and 4) **(p.14)**

**Table S7.** Gene ontology enrichments (TOP 100) for enhancers with an activity shift in the guinea pig branch (related to Figure 3) (related to Figures 3) **(p.15)**

**Table S8.** Transcription factor binding site enrichments for enhancers with an activity shift in the ancestral mole-rat branch (related to Figures 3) **(p.15)**

**Table S9.** Significant association between enriched transcription factor binding sites and ontology terms for enhancers with an activity shift in the ancestral mole-rat branch (related to Figures 3) **(p.15)**

**Table S10.** Clustering of enriched gene ontologies in ancestral and single-species mole-rat branches (related to Figure 4). **(p.15)**

**Table S11.** Transcription factor binding sites associated with specific repetitive elements in non-alignable mole-rat enhancers (related to Figure 5) **(p.15)**

**Supplemental datasets**:

The following datasets have been deposited to Zenodo (https://doi.org/10.5281/zenodo.7442105), along with the code, inputs and environment to reproduce the presented analyses (prediction of regulatory elements from histone marks peaks, identification of orthologous regions, normalization of reads densities, phylogenetic modeling of regulatory activity shifts and functional enrichment tests for mole-rats regulatory elements).

**Dataset S1:** Promoters, enhancers and primed enhancers predicted in each species before cross-mapping (.bed files).

**Dataset S2:** Orthologous promoters, enhancers and primed enhancers given in each species genomic coordinates system (.bed files), and normalized H3K27ac reads densities at orthologous promoters and enhancers.

**Dataset S3:** UP and DOWN promoters and enhancers in mole-rats (.bed files), and regions to genes association tables.

**Dataset S4:** Non-alignable promoters and enhancers in each species (.bed files), *de novo* repeats annotation for mole-rats (.bed and consensus .fasta files), tables with non-alignable mole-rats enhancers enriched in specific repeats.

### SUPPLEMENTAL FIGURES

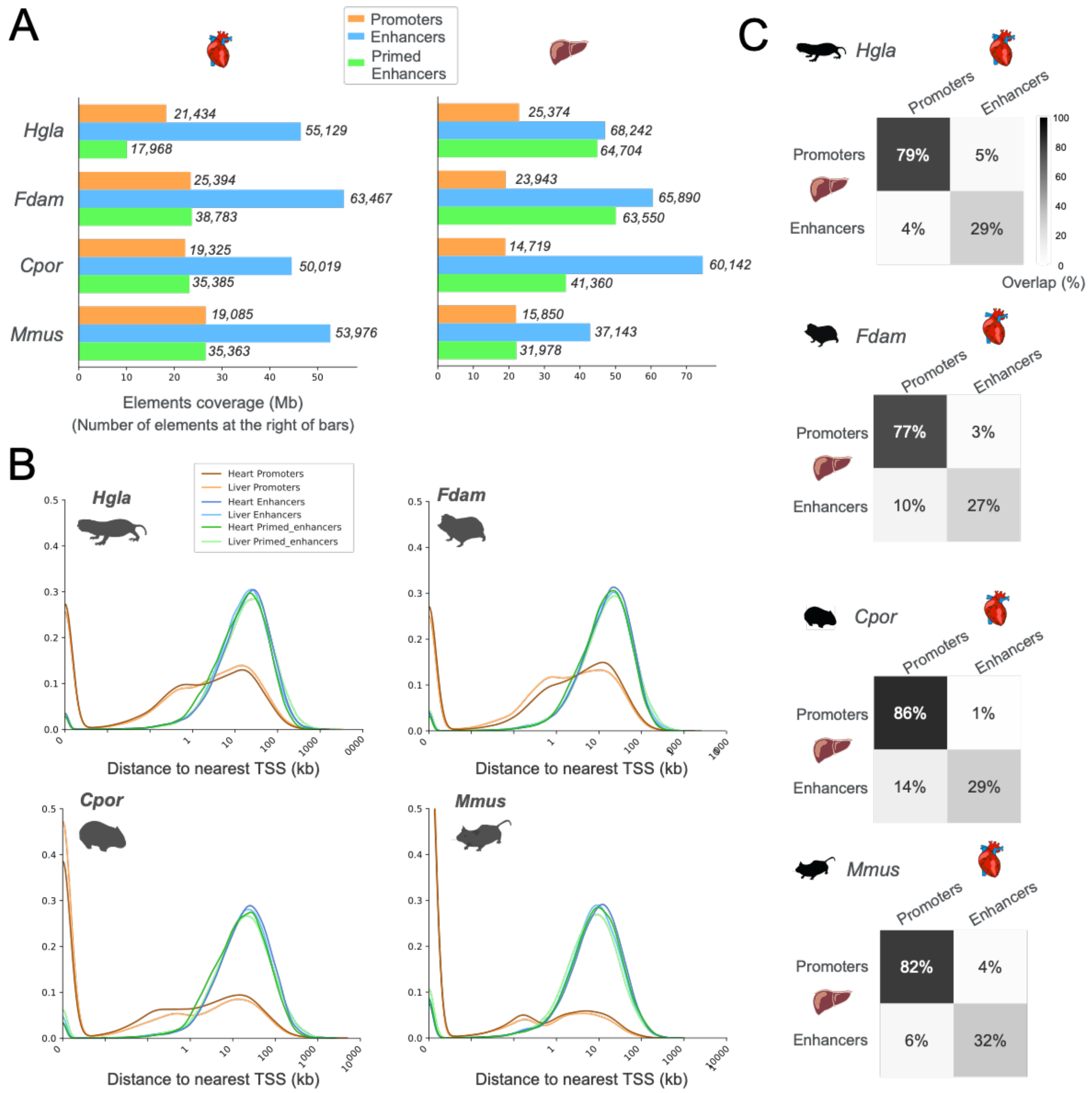


**Figure S1. Basic properties of promoters, enhancers and primed enhancers identified in heart and liver tissues across two mole-rats and two outgroup rodents** (related to Figure 1)

**A.** Number of promoters, enhancers and primed enhancers identified in each species. Bars correspond to total genomic coverage of each set of elements. Number of elements are indicated at the right of bars.

**B.** Distribution of distances to the nearest transcription start site (TSS) for promoters, enhancers and primed enhancers across the two tissues and four species. As expected, identified promoters are typically close to annotated genes, while enhancers are distal regulatory elements.

**C.** Overlap between promoters and enhancers across liver and heart tissues, in each of the four species.

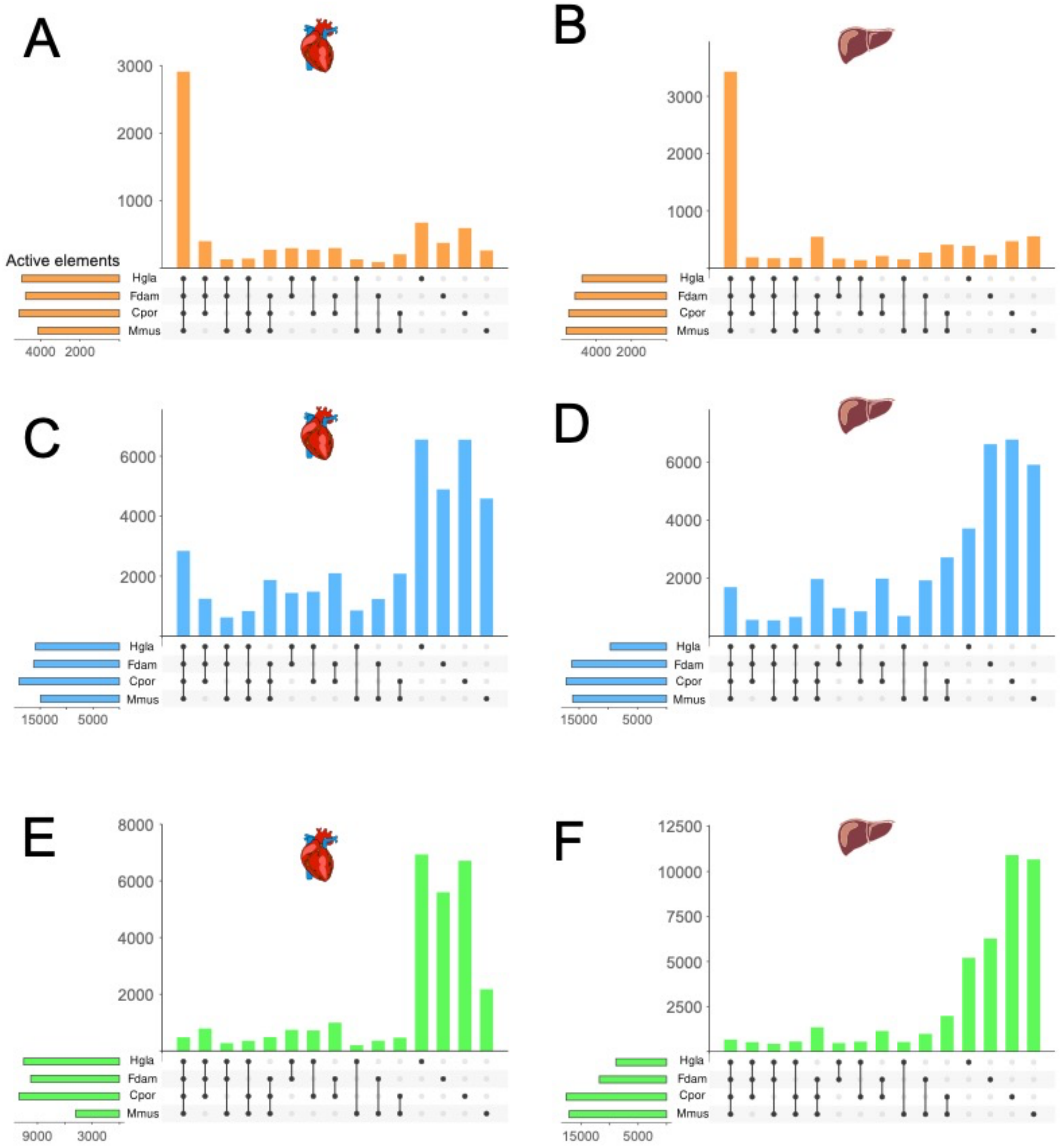


**Figure S2. Cross-mapping of promoters, enhancers and primed enhancers across the four study species** (related to Figure 1)

**A-F.** Activity of orthologous promoters, enhancers and primed enhancers across species. Upset plots indicate the number of orthologous promoters (orange), enhancers (blue) and primed enhancers (green) active across several species or species-specific. In agreement with previous observations, promoter epigenomic activity is highly conserved across species while enhancers and primed enhancers are mostly species-specific.

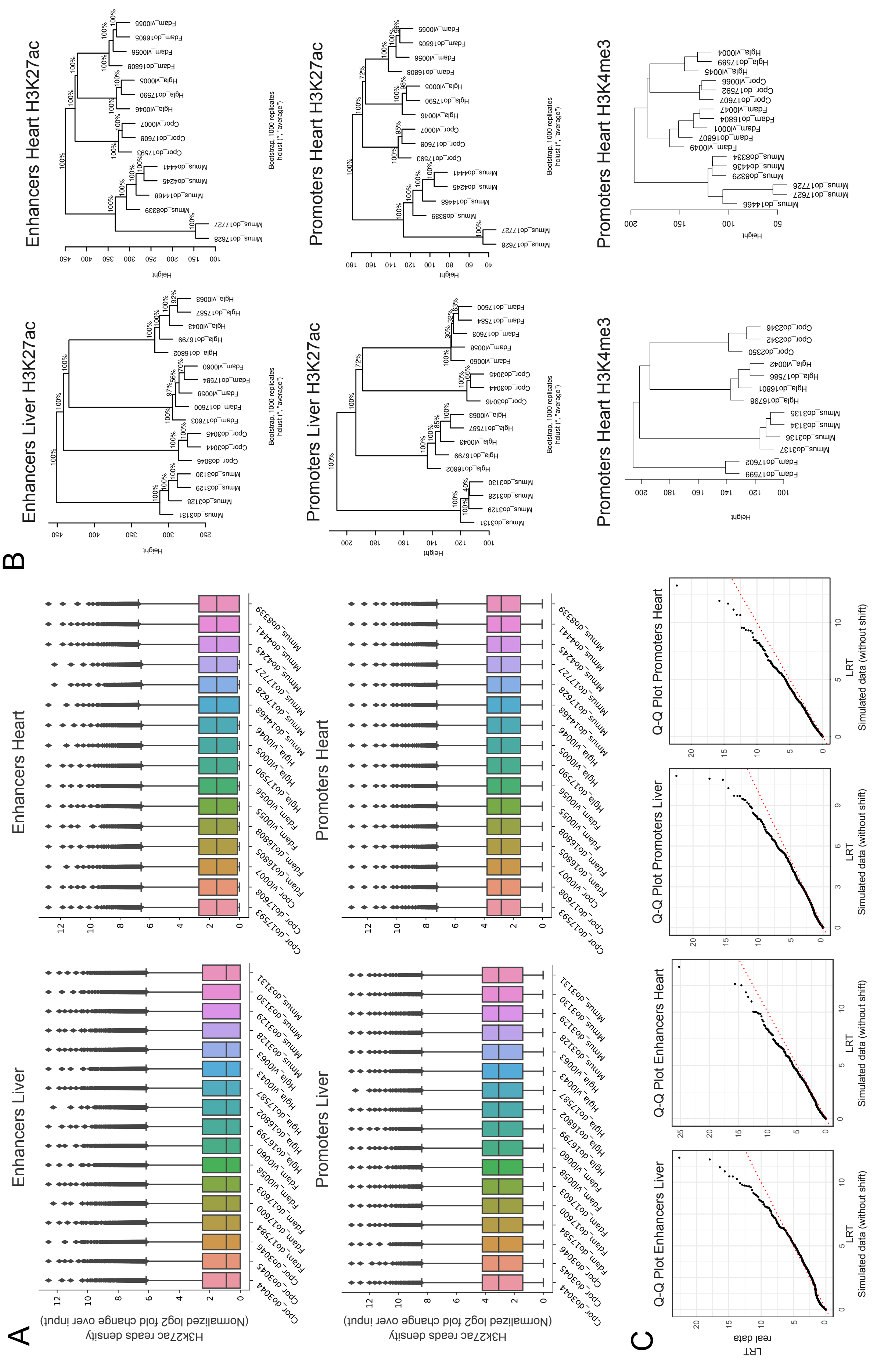
A
Enhancers Liver
Enhancers Heart
Promoters Liver
Promoters Heart
H3K27ac reads density
(Normalized log2 fold change over input)
B
Enhancers Liver H3K27ac
Enhancers Heart H3K27ac
Promoters Liver H3K27ac
Promoters Heart H3K27ac
Promoters Heart H3K4me3
Promoters Heart H3K4me3
Bootstrap, 1000 replicates
hclust (*, "average")
Height
C
Q-Q Plot Enhancers Liver
Q-Q Plot Enhancers Heart
Q-Q Plot Promoters Liver
Q-Q Plot Promoters Heart
LRT
real data
LRT
Simulated data (without shift)

**Figure S3. Data normalisation and parameter estimation for phylogenetic modeling** (related to Figure 2)

**A.** Distribution of normalized H3K27 reads densities at orthologous enhancers and promoters. Reads densities were expressed as log2 fold-change of signal over input ChIP and normalized with quantile normalization.

**B.** Hierarchical clustering of normalized reads densities at orthologous enhancers and promoters. Hierarchical clustering was performed using the euclidean distance and average linkage, with bootstrap support computed from resampling using the shipunov R package. Clustering with normalized H3K27ac read densities recapitulates the species phylogeny with high support. Conversely, H3K4me3 normalized read densities lack phylogenetic signal and are not suitable for phylogenetic modeling with EVE.

**C.** Q-Q plots comparing distributions of Likelihood Ratio Tests (LRT) statistics from the EVE branch-shift test. LRT were computed from data simulated under the null (no shift of regulatory activity, x-axis) and observed data (y-axis). LRTs from observed and simulated data follow the same distribution for low LRT values with a departure from the null for observed data at high LRT values, showing that simulated LRTs can be used to compute empirical p-values.

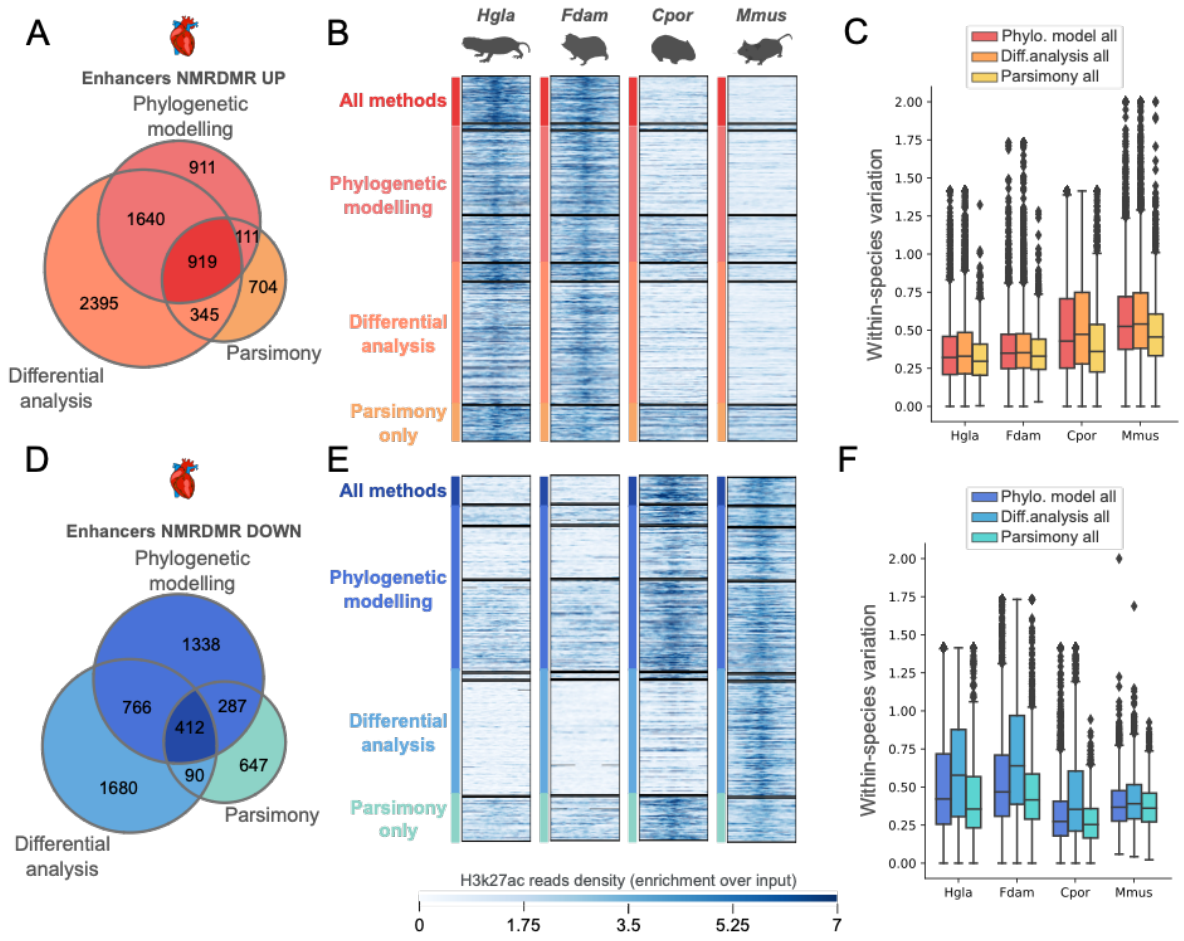


**Figure S4. Comparison of phylogenetic modeling results in heart with regulatory shifts inferred from parsimony and differential binding analysis** (related to Figure 2)

**A-C.** Comparison of heart enhancers identified by phylogenetic modeling, differential analysis and parsimony as upregulated in both mole-rats (Up enhancers, ancestral branch) **A.** Venn diagram showing the overlap between elements identified by each method. **B.** Read density heatmaps for elements identified by the different approaches; black lines indicate overlapping subsets across approaches (as denoted in A.). H3K27ac read density is presented as fold enrichment over input, averaged across biological replicates. Colors on y-axis correspond to category combinations highlighted on the Venn diagram in A. **C.** Within-species variation (coefficient of variation) in read density for enhancers identified with the different methods. Up elements identified with differential analysis show systematically higher variation, suggesting they could contain a higher fraction of false positives.
**D-F.** Comparison of heart enhancers identified by phylogenetic modeling, differential analysis and parsimony, as downregulated in both mole-rats (Down enhancers, ancestral branch). Representation as in A-C.

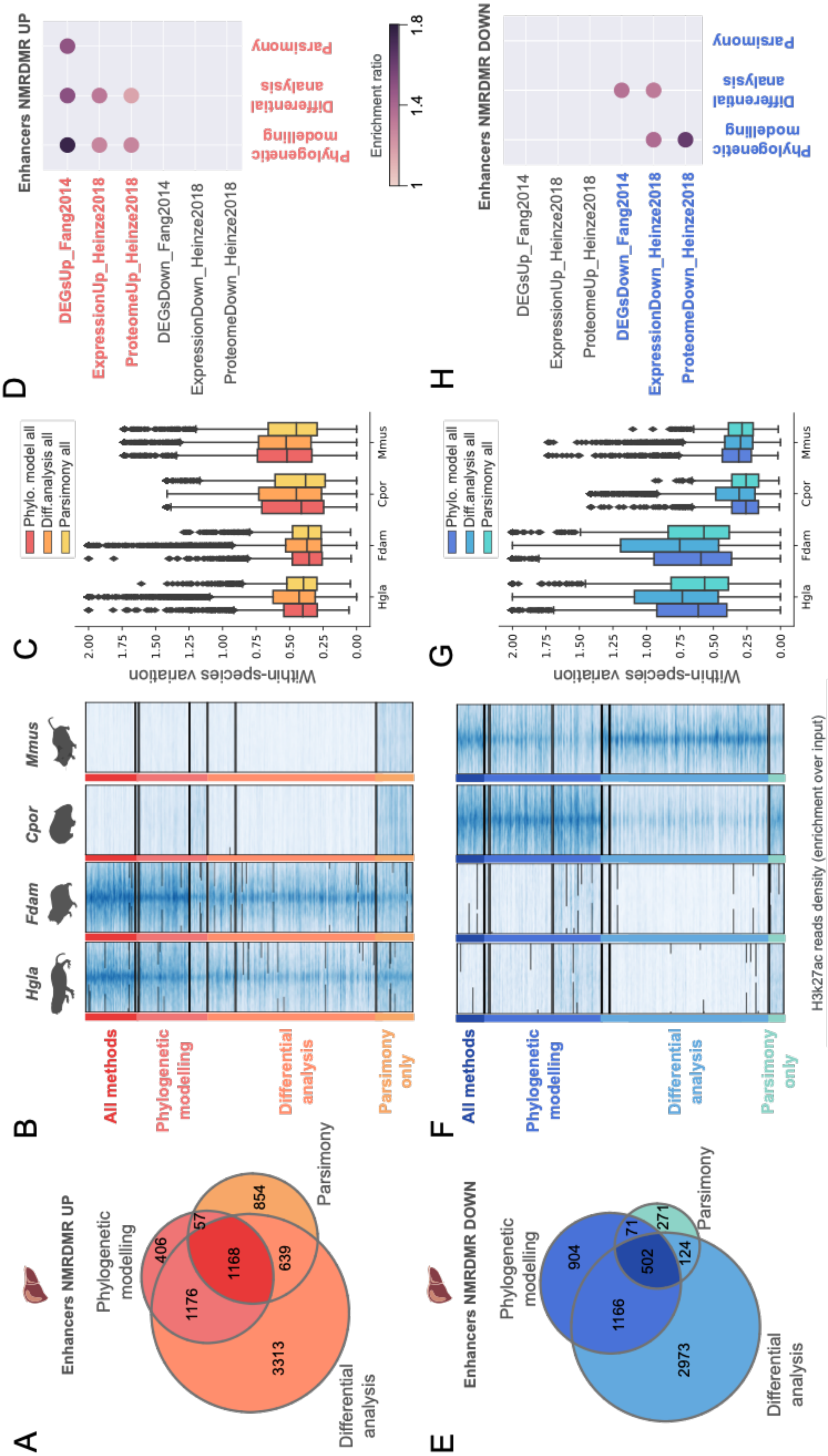
A
Enhancers NMRDMR UP
Phylogenetic modelling
Parsimony
Differential analysis
406
57
854
1168
639
1176
3313
B
All methods
Phylogenetic modelling
Differential analysis
Parsimony only
Hgla
Fdam
Cpor
Mmus
C
Within-species variation
Phylo. model all
Diff.analysis all
Parsimony all
D
Enhancers NMRDMR UP
DEGsUp_Fang2014
ExpressionUp_Heinze2018
ProteomeUp_Heinze2018
DEGsDown_Fang2014
ExpressionDown_Heinze2018
ProteomeDown_Heinze2018
Phylogenetic modelling
Differential analysis
Parsimony
Enrichment ratio
E
Enhancers NMRDMR DOWN
Phylogenetic modelling
Parsimony
Differential analysis
904
71
271
502
124
1166
2973
F
All methods
Phylogenetic modelling
Differential analysis
Parsimony only
H3k27ac reads density (enrichment over input)
G
Within-species variation
Phylo. model all
Diff.analysis all
Parsimony all
H
Enhancers NMRDMR DOWN
DEGsUp_Fang2014
ExpressionUp_Heinze2018
ProteomeUp_Heinze2018
DEGsDown_Fang2014
ExpressionDown_Heinze2018
ProteomeDown_Heinze2018
Phylogenetic modelling
Differential analysis
Parsimony

**Figure S5. Comparison of phylogenetic modeling results in liver with regulatory shifts inferred from parsimony and differential binding analyses** (related to Figure 2)

**A-D.** Comparison of liver enhancers identified by phylogenetic modeling, differential analysis and parsimony as upregulated in both mole-rats (Up enhancers, ancestral branch). **A-C** as in Fig. S4. **D.** Association of identified Up elements with previously reported differentially-expressed genes (DEGs) in mole rats (see GREAT enrichment tests, Methods). Phylogenetic modeling recovers elements significantly associated with DEGs, and with higher enrichment ratios than other methods.
**E-H.** Comparison of liver enhancers identified by phylogenetic modeling, differential analysis and parsimony, as downregulated in both mole-rats (Down enhancers, ancestral branch). Representation as in A-D.

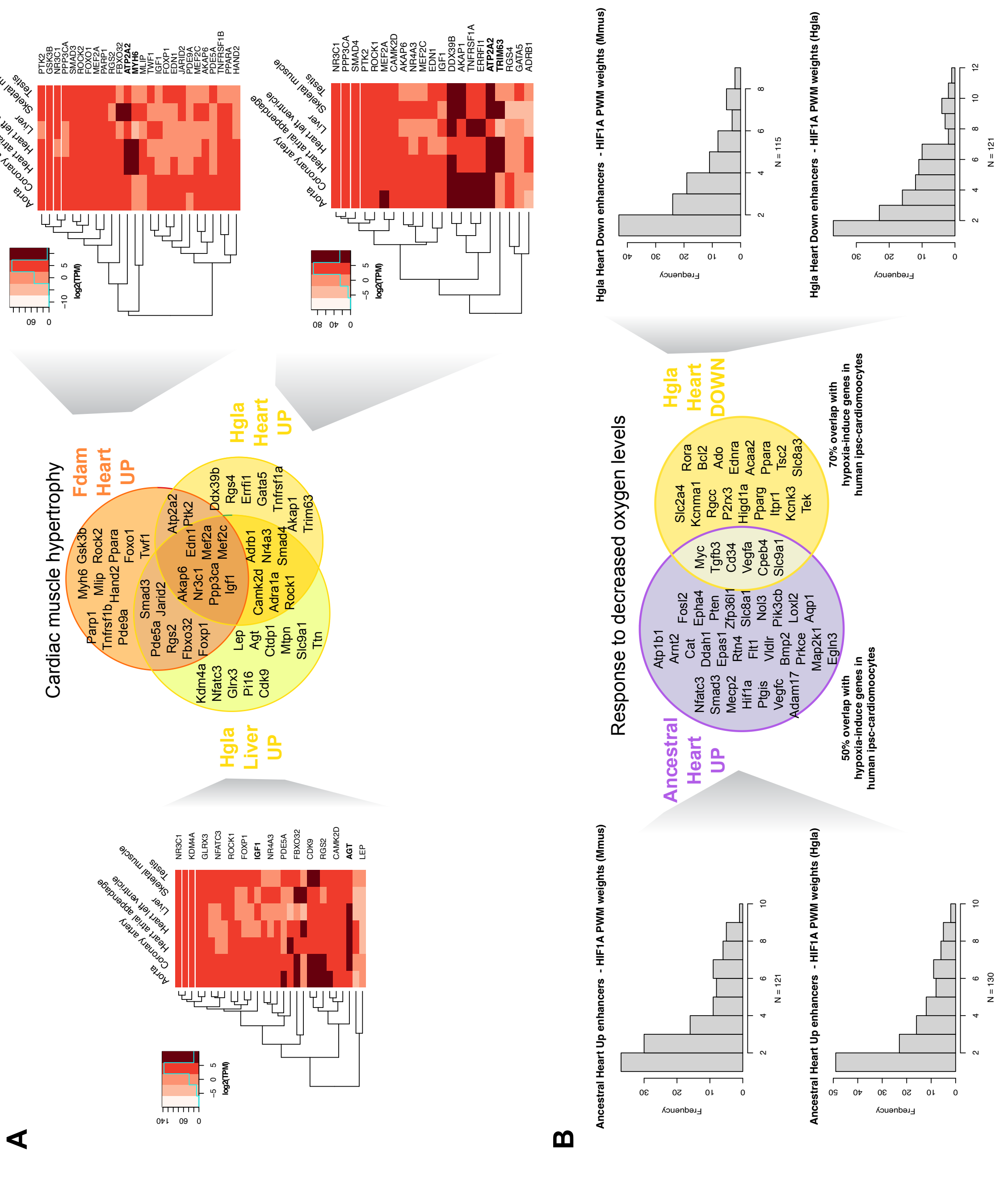


**Figure S6. Additional properties of ancestral and single-species enhancer shifts in selected ontology categories** (related to Figure 4 and Discussion)

**A.** Gene expression heatmaps showing human gene expression levels across selected tissues from GTEX v8 data. Represented genes correspond to those associated to the Cardiac hypertrophy ontology term and Up liver enhancers in the naked mole-rat branch (left), Up heart enhancers in the Damaraland mole-rat branch (upper right), and Up heart enhancers in the naked mole-rat branch (lower right).

**B.** Comparison of HIF binding affinities for mole-rat enhancer shifts proximal to genes in the Response to hypoxia ontology category, specifically for Up heart enhancers in the ancestral mole-rat branch (left) and Down heart enhancers in the naked mole-rat branch (right). For each, histograms show the distribution of predicted HIF binding affinities for orthologous locations of each elements set in mouse (top) and naked mole-rat (bottom), using the HIF1A position-weight matrix from JASPAR database (Core Vertebrate collection). The mouse and naked mole-rat HIF binding affinity score distributions are not significantly different for Up heart enhancers in the ancestral branch (Kolmogorov-Smirnov test, p=0.82). However, for Down heart enhancers in the naked mole-rat branch, predicted HIF binding affinities tend to be higher in mouse orthologous sequences compared to naked mole-rat sequences (Kolmogorov-Smirnov test, p=0.05).

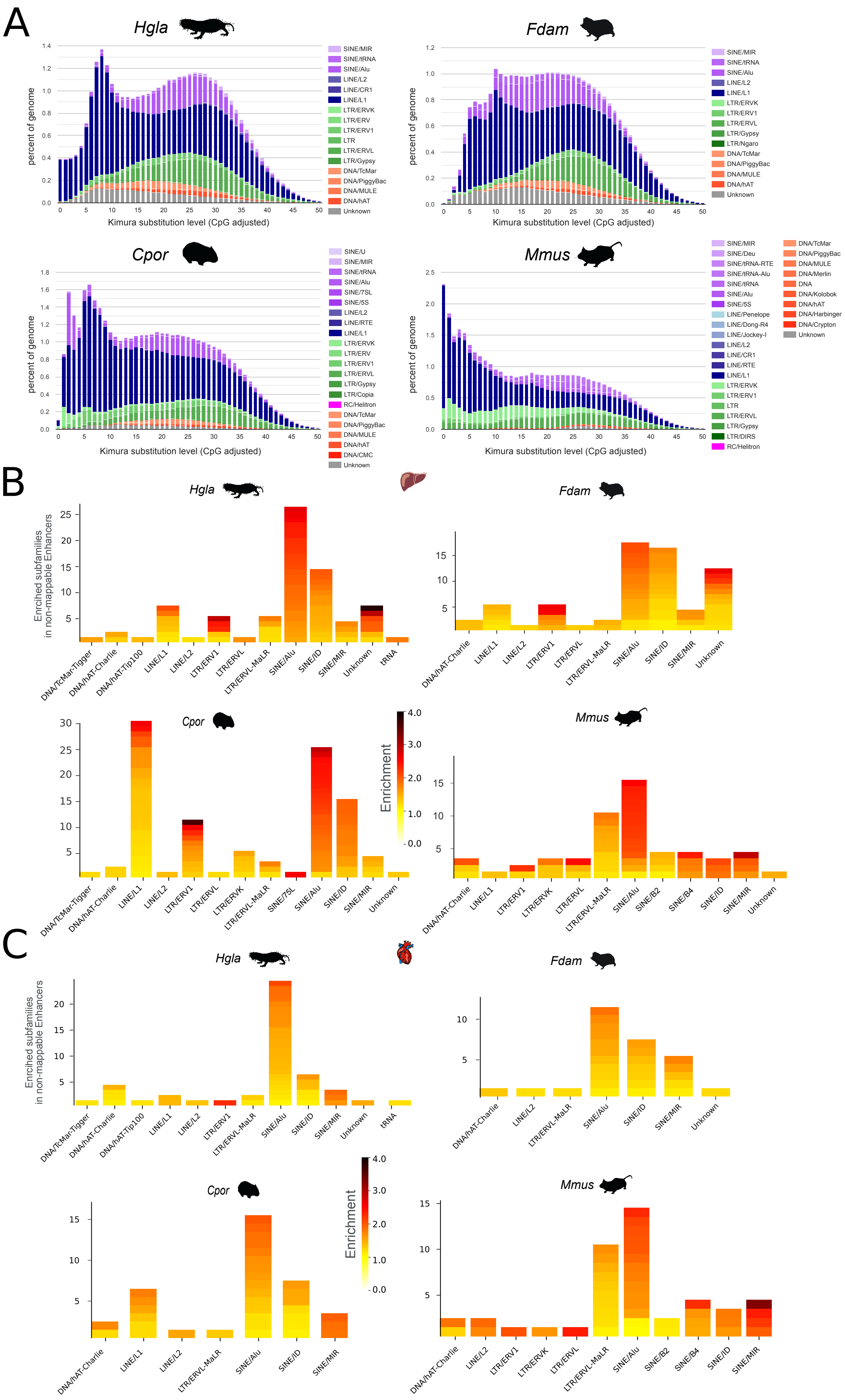
A
Hgla
Fdam
Cpor
Mmus
percent of genome
Kimura substitution level (CpG adjusted)
B
Enriched subfamilies in non-mappable Enhancers
Enrichment
C

**Figure S7. Additional properties of repetitive elements in mole-rat genomes and their association with non-alignable enhancers** (related to Figure 5)

**A.** Landscape plots for repeat annotations in each of the four rodents. For mole-rats, we generated *de novo* repeat libraries with RepeatModeller. For mouse and guinea pig, we re-used publicly available *de novo* repeat libraries from Dfam. Landscape plots were drawn using scripts from the RepeatMasker suite.

**B-C.** Repeat families enriched in non-alignable enhancers in liver (B) and heart (C) for each of the four rodents (permutation tests, Methods, corrected p-values <0.01). For each family, the number of enriched sub-family is indicated by the height of bars, colours for each sub-family are proportional to enrichment ratios.

### SUPPLEMENTAL TABLES

**Table S1**: Species and tissue samples used in this study (related to Methods, Figure 1 and Figure S1) **[tableS1.xlsx file]**

For the four species in the dataset, we provide here experimental details regarding origin of the samples, sex, ages, breeding class and number of replicates for each profiled histone mark and tissue.

**Table S2.** Simulations to calibrate statistical thresholds for the branch-shift test of the phylogenetic modeling approach (related to Figure 2) **[tableS2.xlsx file]**

For the different elements sets and mole-rat branches we performed simulations to assess statistical power of the branch-shift test at different statistical thresholds (Methods). We report here the AUC values, false and true positive rates and false discovery rate at the selected significance threshold (alpha=0.20, absolute value of shift > 1.5), estimated under various simulation settings.

**Table S3.** Genes significantly associated to enhancers with an activity shift in the ancestral mole-rat branch (related to Figure 2) **[tableS3.xlsx file]**

For each element set, we report genes signficantly associated to enhancers with an acitivty shift in the ancestral mole-rat branch (GREAT enrichment tests using the DMR genome, Methods). Gene names were transferred from mouse orthologs using orthologies from ensembl 102. We report the Damaraland mole-rat Ensembl gene ID when no ortholog was found.

**Table S4.** Gene ontology enrichments (TOP 100) for enhancers with an activity shift in the ancestral mole-rat branch (related to Figures 3 and 4) **[tableS4.xlsx file]**

For each element set, we report gene ontologies signficantly associated to enhancers with an activity shift in the ancestral mole-rat branch (GREAT enrichment tests using the DMR genome, Methods).

Columns are as follows; GO: gene ontology id, name: gene ontology name, Binom Enrichment:, enrichment ratio for the binomial test over regions, # foreground regions: number of regions associated with the GO, # expected regions: exepected number of regions associated with the GO, Binom BH p-value: corrected p-value for the binomial test over regions, Hyper Enrichment: enrichment ratio for the hypergeometric test over genes, # foreground genes: number of foreground genes associated with the GO, # expected genes: expected number of genes associated with the GO, Hyper BH p-value: corrected p-value for the hypergeometric test over genes, Genes: foreground genes associated with the GO, Regions: foreground regions associated with the GO.

**Table S5.** C2 pathway enrichments for enhancers with an activity shift in the ancestral mole-rat branch (related to Figures 3 and 4) **[tableS5.xlsx file]**

Columns as in Table S4.

**Table S6.** Gene ontology enrichments for promoters with an activity shift in the ancestral mole-rat branch (related to Figures 3 and 4) (p.21) **[tableS6.xlsx file]**

Columns as in Table S4. Note that no significant enrichment was found for liver promoters.

**Table S7.** Gene ontology enrichments (TOP 100) for enhancers with an activity shift in the guinea pig branch (related to Figure 3) **[tableS7.xlsx file]**

Columns as in Table S4.

**Table S8.** Transcription factor binding site enrichments for enhancers with an activity shift in the ancestral mole-rat branch (related to Figures 3) **[tableS8.xlsx file]**

For each set (Liver enhancers up, Liver enhancers down, Heart enhancers up, Heart enhancers down) columns are as follows: HOMER TFBS name, TFBS consensus, p-value, log(p-value), q-value, number and percentage of target sequences with TFBS, number and percentage of backrgound sequences with TFBS.

**Table S9.** Significant association between enriched transcription factor binding sites and ontology terms for enhancers with an activity shift in the ancestral mole-rat branch (related to Figures 3). **[tableS9.xlsx file]**

Columns are as follows: element set (Liver enhancers up, Liver enhancers down, Heart enhancers up, Heart enhancers down), TFBS – GO association, corrected p-value, Enrichment, Enhancer IDs.

**Table S10.** Clustering of enriched gene ontologies in ancestral and single-species mole-rat branches (related to Figure 4). **[tableS10.xlsx file]**

Columns are as follows: Gene ontology terns or C2 pathway name, enhancer set(s) in which the term is enriched, name of the cluster.

**Table S11.** Transcription factor binding sites associated with specific repetitive elements in non-alignable mole-rat enhancers (related to Figure 5). **[tableS11.xlsx file]**

This file contains: enriched TFBS – repeats pairs in mole-rats, enriched RAR – and HNF4 – repeats pairs in outgroups, enriched gene ontologies in mole-rats HNF4 – SINE/ID and cluster memberships for GO enriched in RAR – SINE/Alu in each of the 4 species.
